# Sex-Dependent Effects of Chronic Microdrive Implantation on Acquisition of Trace Eyeblink Conditioning

**DOI:** 10.1101/2021.07.30.454487

**Authors:** Amy P Rapp, Timothy J Hark, John M Power, Jeffrey N Savas, M Matthew Oh, John F Disterhoft

## Abstract

Neuroscience techniques, including in vivo recording, have allowed for a great expansion in knowledge; however, this technology may also affect the very phenomena researchers set out to investigate. Including both female and male mice in our associative learning experiments shed light on sex differences on the impact of chronic implantation of tetrodes on learning. While previous research showed intact female mice acquired trace eyeblink conditioning faster than male and ovariectomized females, implantation of chronic microdrive arrays showed sexually dimorphic effects on learning. Microdrive implanted male mice acquired the associative learning paradigm faster than both intact and ovariectomized females. These effects were not due to the weight of the drive alone, as there were no significant sex-differences in learning of animals that received “dummy drive” implants without tetrodes lowered into the brain. Tandem mass tag mass spectrometry and western blot analysis suggest that significant alterations in the MAPK pathway, acute inflammation, and brain derived neurotrophic factor may underlie these observed sex- and surgery-dependent effects on learning.

## Introduction

Electrophysiology has allowed neuroscientists to study a variety of phenomena including the activity of individual neurons and larger population encoding of groups of neurons have been studied with in vivo neuronal recording techniques (Obien et al. 2014). In fact, many fundamental neuroscience advances have come from extracellular recordings over the last half a century, including: place cells (O’Keefe and Dostrovsky 1971), orientation-selective cells (Hubel and Wiesel 1959), grid cells (Hafting et al. 2005), and time cells (Manns, Howard, and Eichenbaum 2007). In-vivo recording remains a critical tool for neuroscience research, especially as it expands with new technologies (Hong and Lieber 2019). Therefore, it is necessary to understand how the implementation of chronic recordings affects acquisition of learning and memory.

Stress exposure can activate the hypothalamic-pituitary-adrenal (HPA) axis, leading to production of glucocorticoids, neurotransmitters and neuropeptides (Lindau, Almkvist, and Mohammed 2016). Glucocorticoids alter many brain regions, including the hippocampus, which contains a large number of glucocorticoid receptors (Moreira et al. 2016). As a result, chronic stress can cause impaired hippocampal-dependent function (Moreira et al. 2016) through several mechanisms such as an increase in neurodegeneration and decrease in neurogenesis (McEwen 2000; Pham et al. 2003), an increase in glutamate (Karst et al. 2005; Magariños, García Verdugo, and Mcewen 1997; Wiegert, Joëls, and Krugers 2007), a decrease in Brain Derived Nerve Growth Factor (NGF) (Bath, Schilit, and Lee 2013; Lakshminarasimhan and Chattarji 2012; Smith and Cizza 1996), and a retraction of dendrites (Gould et al. 1997; Magarinños and McEwen 1995; Sousa et al. 2000). Notably, stress exposure before learning can either enhance (Domes et al. 2002; Smeets et al. 2007; Weiss et al. 2005) or impair acquisition (Diamond et al. 2006; Elzinga and Roelofs 2005; Kirschbaum et al. 1996; Schwabe et al. 2012). Sexually dimorphic effects of stress on learning have been shown on a variety of behavioral tasks. In response to stress from a repeated tail shock, male rats acquired trace eyeblink conditioning (tEBC) faster than their female and unstressed male counterparts (Shors et al. 1998; Wood and Shors 1998). Similarly, in response to high-dose injections of corticosterone, eyeblink conditioning acquisition was facilitated in male Long-Evans rats, but there was no impact on acquisition in female rats (Wentworth-Eidsaune, Hennessy, and Claflin 2016). Corticosterone treatment also enhanced learning in male rats on an auditory fear conditioning task, but impaired learning in female rats (Lesuis et al. 2018). These sexually dimorphic behavioral responses to stress suggest that stress may underlie the behavioral differences initially observed in chronically implanted animals.

We set out to investigate if the weight of a microdrive implant into the brain acts as a chronic stressor to mice, thereby affecting the learning rate of a hippocampus-dependent task in a sex dependent manner. Mice with chronically implanted microdrive arrays were compared to those with a “dummy drive” (no tetrodes implanted into the brain) as well as to those with smaller and lighter headbolt implants that are used for eyeblink conditioning. We found that compared to headbolt implanted control mice, implantation of a chronic microdrive array with tetrodes lowered into the cortex facilitated tEBC acquisition in male mice, while both microdrive and “dummy drive” implantation tended to impair tEBC acquisition in intact female mice. Tandem mass tag mass spectrometry (TMT-MS) and western blot analysis suggest that significant alterations in the MAPK pathway, acute inflammation, and brain derived neurotrophic factor (BDNF) may underlie sex and surgery dependent effects in acquisition. These findings also emphasize the need for inclusion of both sexes in research, in order to evaluate possible sex-specific effects that neuroscience technologies and methods may produce.

## Methods

### Animals

All procedures were approved by and completed in accordance with the Northwestern University Animal Care and Use Committee guidelines. Experiments were performed with young adult (3-4 months) male, intact female and ovariectomized female (ovx) C57BL/6J mice. All mice were obtained from Jackson Laboratory (Bar Harbor, Maine). Ovariectomy was performed by Jackson Laboratory at least two weeks prior to shipment. Estrous cycles of female mice were not monitored as previous studies have demonstrated females without a staged estrous cycle had similar variability as males in behavioral tasks (Prendergast, Onishi, and Zucker 2014; Rapp et al. 2021). All mice were housed in Northwestern University temperature-controlled facilities in a 14-hour light: 10-hour dark cycle and fed ad lib. Mice were group housed at arrival and allowed to acclimate to Northwestern University facilities for at least one week prior to experimentation. After implantation surgery, mice were housed individually.

### Surgery

#### Headbolt Surgery

Headbolt animals were previously utilized for Rapp et al 2021. Male (n=18), intact female (n=18) and ovariectomized female (n=14) mice were implanted with a custom headbolt two weeks prior to behavioral training. Animals were briefly anesthetized with 3-4% vaporized isoflurane mixed with oxygen (flow rate: 1-2 liters/minute). Buprenorphine (0.05-2 mg/kg) was administered subcutaneously as an analgesic. The scalp was shaved, and the mouse was placed in a stereotaxic device. The scalp was sterilized with iodine and 70% ethanol, then an incision was made along the midline with a scalpel blade (No 15). The skin was retracted laterally with microclips, and the skull was cleaned with 3% hydrogen peroxide then sterile saline. The skull was scored with the scalpel blade to promote adhesion of the dental cement. Two small stainless-steel screws (00-90) were implanted to the left of midline (one in front of Bregma, and one in front of Lambda). The bare stainless steel groundwire (0.005in:AM Systems: 792800) of the custom headbolt was wrapped around the screws in a figure-eight pattern to serve as a ground for EMG recordings. A thin layer of Metabond adhesive cement was spread over the skull, screws, and wire to secure them in place. To expose the muscle and place EMG wires for EMG recording, the skin surrounding the right eye was retracted. Four polyimide-coated stainless steel (0.005in: PlasticsOne: 005sw/2.0 37365 SS) wires with 2-3 mm of exposed wire were placed on the muscularis orbicularis oculi. The headbolt piece and base of the EMG recording wires were then secured with additional adhesive cement. The skin was released from microclips and placed over the cement. Skin was allowed to rest naturally, and the exposed area was sealed with additional adhesive cement. Animals recovered on a warm heating pad before being returned to their home cage. Animals were allowed five to seven days to recover before habituation began.

#### Dummy Drive Implant Surgery

Animals in the Dummy Drive group (Males n=18, Intact Females n=17, Ovariectomized Females n= 6) were implanted with a modified headbolt containing one coated stainless steel groundwire (0.0110 in coated: AM Systems: 79200) and two polyimide-coated stainless steel (0.005in: PlasticsOne: 005sw/2.0 37365 SS) wires similar to the process previously described. Dummy Drive animals were also implanted with a Neuralynx Halo-10-Mini microdrive with either a Quick Clip or Omnetics electronic interface board (EIB) connector (Neuralynx, Bozeman, MT). The exit tip of the Halo-10-Mini microdrive was covered with silicone lubricant (Danco 88693P) and the microdrive was cemented to the skull with Metabond dental adhesive (Parkell Inc.) and Hygenic dental cement (Patterson Dental). After the microdrive was secured and the skin was placed and cemented in its natural state, 3D printed head-fixation bars were cemented perpendicular to the skull, above the ears. Animals recovered on a warm heating pad before being returned to their home cage. Animals were allowed five to seven days to recover before habituation began. Chow was placed in a glass bowl (Amazon, B08KNTWCDD) at the bottom of the cage as the wire-top was removed to prevent damage to the implant. After surgery, mice were housed individually.

#### Microdrive Implant Surgery

Microdrive implant animals (Males n= 13, Intact Females n=17, Ovariectomized Females n= 9) were implanted with the same modified headbolt as the Dummy Drive group plus a Omnetics connector Custom 7-Degree Neuralynx Halo-10-Mini Microdrive (n= 27) or custom 3D-printed microdrive (n= 12). Prior to implantation of the modified headbolt, the skull (bregma-lambda) was leveled. Two stainless-steel skull screws were implanted on either side of the coronal suture (0-80 Screw, 91772a049, McMaster Carr) for the headbolt ground. An additional stainless steel skull screw (00-90) was implanted above the cerebellum for the microdrive ground wire. A craniotomy was made (Drill bit: Stoeling Co: 0.45mm: 514551) at AP: +3.3 ML: -3.2. The exit tip of the microdrive was coated in silicone lubricant and the microdrive was lowered into place. The tips of the tungsten tetrodes extended past the exit tip of the microdrive and were lowered marginally into the cortex at implantation. The microdrive was cemented in place with multiple coats of Metabond dental adhesive and dental cement. After the skin was placed naturally around the dental cement and secured, 3D printed head-fixation bars were cemented in place as well. Animals recovered on a warm heating pad before being returned to their home cage. Chow was placed in a glass bowl (Amazon, B08KNTWCDD) at the bottom of the cage as the wire-top was removed to prevent damage to the implant. After surgery, mice were singly housed.

### Trace Eyeblink Conditioning

#### Headbolt

Procedures for conditioning for headbolt implanted animals was previously described in (Rapp et al. 2021). Prior to behavioral training, mice were handled for three days for five minutes/day to habituate mice to restraint and the experimenter. After three days of handling, mice were habituated to head-fixation on a moveable cylinder apparatus for the length of a training session without the presentation of stimuli. Training began two days following habituation. Training consisted of one session per day for ten days (Figure 1b). Conditioned animals received a 65±2 dB tone (250ms, 2kHz) conditioned stimulus (CS) paired with a 35±5 PSI corneal airpuff (30-40 ms) unconditioned stimulus (US) (Figure 1b). The CS and US were separated by a 250 ms trace interval in which no stimuli were presented. Each conditioning session consisted of 50 paired CS/US trials with a random 35-55 second inter-trial interval (Figure 1b). Custom routines in LabVIEW (National Instruments) were used for stimulus presentation procedures, data collection, storage, and analysis (routines available upon request). Tone intensity was calibrated with a sound meter, placed where the mouse would be, at the start of each day of training. Air pressure was calibrated with a manometer (Thermo Fisher Scientific) secured at the output of a 0.5-inch 16-gauge blunted needle before each training session. Animals were visually monitored during training through a camera (Logitech C270) attached to the frame of the cylinder apparatus (Figure 1d). Trials were not presented when the animal was visibly moving and as detected by the behavioral control software.

**Figure 1.**
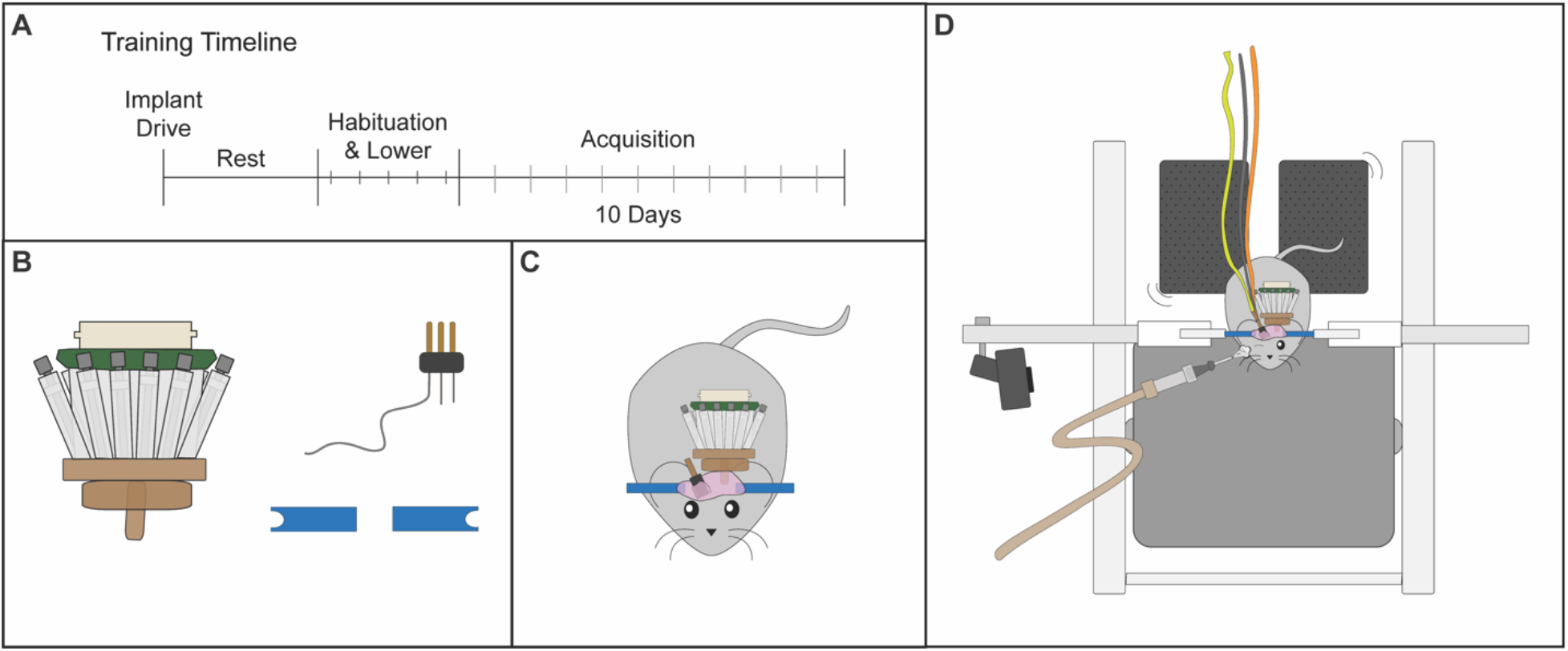
A. Training Timeline. B. Microdrive implant, headbolt, earbars for fixation C. Implant depicted on mouse. D. Behavioral training setup depicting mouse running atop cylinder.

#### Microdrive and Dummy Drive

Prior to behavioral training, mice were habituated to the head-fixed apparatus for forty minutes/day for five days. During these habituation sessions, tetrodes were lowered into the Entorhinal Cortex for microdrive animals (Figure 1a). Eyeblink training consisted of one session per day for ten days (Figure 1a). Conditioned animals received a 65±2 dB tone (250ms, 2kHz) conditioned stimulus (CS) paired with a 35±5 PSI corneal airpuff (30-40 ms) unconditioned stimulus (US) separated by a 250 ms trace interval. Each conditioning session consisted of 50 paired CS/US trials with a random 35-55 second inter-trial interval. Stimulus presentation and calibration was performed similarly to the headbolt animal group. Trials were automatically paused by the LABVIEW software when EMG baseline exceeded 0.25V. Trials were restarted when the EMG baseline was below 0.25V for two consecutive seconds. Animals were visually monitored during training through a camera (Logitech C270) attached to the frame of the cylinder apparatus (Figure 1d).

### Data Analysis

EMG signal output was amplified (x5,000) and filtered (100Hz-5kHz), then digitized at 3kHz and stored by computer. For analysis, EMG data were rectified and integrated with a 10ms time constant. A conditioned response (CR) was defined as increased EMG activity lasting at least 15ms with an amplitude at least 4 standard deviations above the mean baseline activity. Baseline activity was the average EMG activity starting 250ms before CS onset. Trials were excluded if baseline activity was 2 standard deviations above the mean baseline activity for the session. CR onset was calculated in reference to the start of the tone CS. An adaptive CR was defined as a CR that was present in the 200ms before US onset. Animals that reached at least 60% adaptive CRs were considered to have learned the task. The number of trials to 8 consecutive CRs was also used as a measure of learning.

Data were analyzed with Two-Way ANOVA or Mixed-Effects Analysis, One-Way ANOVA, and post-hoc Tukey’s Multiple comparisons test, when appropriate (Prism v8) (Table 1). The probability level of p< 0.05 was used as an indicator of statistical significance. Data are expressed with standard error of the mean. Statistical tests did not include data from habituation, except for direct habituation comparison. Mice were excluded from analysis due to poor health, poor EMG signal, or failure to learn delay conditioning (intact female n=2; male n=4; ovx=1). Delay conditioning is non-hippocampal dependent, where the stimuli overlap. Failure to learn delay conditioning indicates a possible brainstem/cerebellar deficit (Cheng et al., 2008; Heiney et al., 2014; Yang et al., 2015).

### Corticosterone Measurement

To measure stress with a noninvasive technique, corticosterone was measured from fecal samples. Compared to collecting blood samples from the tail vein or orbital sinus, fecal samples can be collected frequently without causing stress from handling, restraint and sample collection (Touma, Palme, and Sachser 2004). Fecal samples also measure circulating hormone levels over a longer term compared to the short time course of blood samples. This technique allowed us to compare stress levels across groups without directly interfering with the animals and introducing further stress.

Fecal matter was collected from the cages of mice prior to surgery, following habituation week, training week 1 (session 1-5) and training week 2 (sessions 6-10). The sample was stored in 2ml Eppendorf tubes in the -80C freezer following collection. Corticosterone was measured through DetectX Corticosterone Enzyme Immunoassay Kit (Arbor Assay: K014). Corticosterone was extracted according to Arbor Assay’s Steroid Solid Extraction Protocol. Fecal matter was dried and powdered with a mortar and pestle (DOT Scientific: JMD050). 0.2 grams of powdered sample was weighed out and 1mL of ethanol was added per 0.1 gm of solid and sealed in a 2ml tube. Samples were vigorously shaken with a vortex for 30 minutes. Samples were centrifuged at 5,000 rpm at 4°C for 15 minutes. The supernatant was collected into a clean tube, then dried down with vacuum centrifugation and resuspended in 100 µl of ethanol. Samples were diluted 1:20 with Assay Buffer then run on the DetectX Corticosterone Enzyme Immunoassay Kit in the 50 μL format according to the Arbor Assay Protocol. Optical density of the Enzyme Linked Immunosorbent Assay (ELISA) was measured at 450 nm using a Synergy HTX multi-mode microplate reader (Biotek) and compared to a standard curve to determine the final concentration. Results were calculated through the online tool MyAssays (MyAssays Ltd). Corticosterone levels were compared through Two-Way ANOVA (Prism V8).

### Tandem Mass Tag Spectrometry

#### TMT-MS Sample Preparation

TMT-MS sample preparation was performed as previously described (Jongkamonwiwat et al. 2020; Rao and Savas 2021). In brief, hippocampal brain extracts were homogenized and protein concentration was determined by BCA assay (Thermo Scientific, Cat# 23225) per manufacturer’s instructions. 300 ug of protein was precipitated with methanol/chloroform to separate proteins from lipids and impurities. The extracted protein was resuspended in 6M guanidine in 100 mM Triethylammonium bicarbonate (TEAB). The proteins were further processed via the reduction of disulfide bonds with dithiothreitol (DTT) and alkylation of cysteine residues with iodoacetamide (IAA). Proteins were then digested with LysC at 1ug/100ug protein (Promega) for 3 h at room temperature (RT) and then digested overnight at 37 °C with Trypsin at 1ug/100ug protein. Following digestion, samples were acidified and desalted using C18 HyperSep columns (ThermoFisher Scientific) per manufacturer instructions. The eluted peptide solution was dried before resuspension in 100 mM TEAB. Micro-BCA assay was subsequently performed to determine the concentration of peptides. 100 μg of peptide from each sample was then used for isobaric labeling. TMT 10-plex labeling was performed on peptide samples according to the manufacturer’s instructions (ThermoFisher Scientific). After incubating for 2 hr at room temperature, the reaction was quenched with 0.3% (v/v) hydroxylamine. Isobaric labeled samples were then combined 1:1:1:1:1:1:1:1:1:1:1:1 and subsequently desalted with C18 HyperSep columns. The combined isobaric labeled peptide samples were fractionated into eight fractions using high pH reversed-phase columns (Pierce). Peptide solutions were dried, stored at −80 °C, and reconstituted in liquid chromatography–mass spectrometry (LC–MS) loading buffer (2% acetonitrile, 0.05% trifluoroacetic acid) for LC–-MS/MS analysis.

#### TMT-MS Analysis

TMT-MS analysis was performed as previously described (Jongkamonwiwat et al. 2020). In short, samples were resuspended in 20 μL of MS loading buffer (2% acetonitrile, 0.05% formic acid), and micro-BCA was performed. 3 μg of each fraction was loaded for LC–MS analysis via an auto-sampler with a Thermo EASY nLC 100 UPLC pump onto a vented Pepmap100, 75 μm × 2 cm, nanoViper trap column coupled to a nanoViper analytical column (Thermo Scientific) with a stainless-steel emitter tip assembled on the nanospray flex ion source with a spray voltage of 2000 V. Orbitrap Fusion was used to generate MS data. The chromatographic run was performed with a 4 h gradient beginning with 100% buffer A and 0% B and increased to 7% B over 5 min, then to 25% B over 160 min, 36% B over 40 min, 45% B over 10 min, 95% B over 10 min, and held at 95% B for 15 min before terminating the scan. Buffer A contained 2% acetonitrile (ACN) and 0.1% formic acid in H2O, and buffer B contained 99.9% ACN with 0.1% formic acid. Multinotch MS3 method was programmed with the following parameters: ion transfer tube temp = 300 °C, easy-IC internal mass calibration, default charge state = 2, and cycle time = 3 s. MS1 detector was set to orbitrap with 60 K resolution, wide quad isolation, mass range = normal, scan range = 300–1800 m/z, max injection time = 50 ms, AGC target = 6 × 105, microscans = 1, RF lens = 60%, without source fragmentation, and datatype = positive and centroid. Monoisotopic precursor selection was set to include charge states 2–7 and reject unassigned. Dynamic exclusion was allowed; n = 1 exclusion for 60 s with 10 ppm tolerance for high and low. The intensity threshold was set to 5 × 103. Precursor selection decision = most intense, top speed, 3 s. MS2 settings include isolation window = 0.7, scan range = auto normal, collision energy = 35% CID, scan rate = turbo, max injection time = 50 ms, AGC target = 6 × 105, and Q = 0.25. In MS3, the top 10 precursor peptides selected for analysis were then fragmented using 65% higher-energy collisional dissociation before orbitrap detection. A precursor selection range of 400–1200 m/z was chosen with mass range tolerance. An exclusion mass width was set to 18 ppm on the low and 5 ppm on the high. Isobaric tag loss exclusion was set to TMT reagent. Additional MS3 settings include an isolation window = 2, orbitrap resolution = 60 K, scan range = 120–500 m/z, AGC target = 6 × 105, max injection time = 120 ms, microscans = 1, and datatype = profile.

#### TMT-MS Data Analysis and Quantification

TMT-MS data analysis was performed as previously described (Jongkamonwiwat et al. 2020). In short, protein identification, TMT quantification, and analysis were performed with The Integrated Proteomics Pipeline-IP2 (Integrated Proteomics Applications, Inc., http://www.integratedproteomics.com/). Proteomic results were analyzed with ProLuCID, DTASelect2, Census, and QuantCompare. MS1, MS2, and MS3 spectrum raw files were extracted using RawExtract 1.9.9 software (http://fields.scripps.edu/downloads.php). Pooled spectral files from all eight fractions for each sample were then searched against the Uniprot mouse protein database and matched to sequences using the ProLuCID/SEQUEST algorithm (ProLuCID ver. 3.1) with 50 ppm peptide mass tolerance for precursor ions and 600 ppm for fragment ions. Fully and half-tryptic peptide candidates were included in the search space, all that fell within the mass tolerance window with no miscleavage constraint, assembled, and filtered with DTASelect2 (ver. 2.1.3) through the Integrated Proteomics Pipeline (IP2 v.5.0.1, Integrated Proteomics Applications, Inc., CA, USA). Static modifications at 57.02146 C and 304.2071 K and N-term were included. The target-decoy strategy was used to verify peptide probabilities and false discovery ratios. A minimum peptide length of five was set for the process of each protein identification, and each dataset included a 1% FDR rate at the protein level based on the target-decoy strategy. Isobaric labeling analysis was established with Census 2 as previously described. TMT channels were normalized by dividing it over the sum of all channels. No intensity threshold was applied. The fold change was then calculated as the mean of the experimental group standardized values, and *p*-values were then calculated by Student’s *t*-test.

#### Online Databases for PANTHER

Protein ontologies were determined with protein analysis through the evolutionary relationship (PANTHER) system (http://www.pantherdb.org) in complete biological process categories. The statistical overrepresentation test was calculated by using the significantly different proteins identified from comparing surgical groups for each sex as the query and the aggregated total proteins identified in all three surgical group comparisons as the reference.

### Western Blot

For western blot analysis, mice were transcardially perfused with ice cold 0.1M PBS. For the perfusion, animals were injected with ketamine/xylazine cocktail (91.95mg/ml ketamine, 8.05mg/ml xylazine). The brains were extracted and sagittally hemisected in 0.1 M PBS. One half of the brain was flash frozen in dry ice and ethanol, while the other hemisphere was drop-fixed in 4% PFA. Brain regions were homogenized in homogenization buffer (4 mM HEPES, 0.32 M sucrose, 0.1 mM MgCl2) containing the following protease inhibitors: aprotinin, leupeptin, AEBSF, benzamidine, PMSF, and pepstatin A. A bead based Precellys 24 homogenizer was used. 300 μl of homogenization buffer was used for the hippocampus. Protein concentration was then determined by BCA assay (Thermo Scientific, Cat# 23225) per manufacturer’s instructions, and optical density (OD) at 562 nm was read on a Synergy HTX multi-mode microplate reader (Biotek) and compared with the respective standard curve.

To prepare the samples for Western blot, each sample was mixed with 6 X SDS sample buffer at a 5:1 ratio. The mixtures were boiled at 95°C for 5 minutes. 50 μg of sample was loaded onto a 16% Tris-Glycine polyacrylamide Gel (Thermo Scientific, Cat# XP00165BOX). Gels were run at ∼100 V for ∼2.5 hours, then were wet transferred to a 0.2 μm nitrocellulose membrane. Membranes were subsequently washed with Tris-buffered saline with 0.1% Tween® (TBST) and blocked with Odyssey Blocking Buffer (LI-COR, Cat# 92740003) in PBS for 1 hour then incubated overnight with primary antibody. The next day membranes were washed and incubated in secondary antibody for 1 hour at RT. Blots were imaged on an Odyssey CLx (Li-Cor). The following primary antibodies were used for Western blots: GFAP anti mouse 1:1000 (Thermo Fisher Scientific Cat# MA5-12023, RRID:AB_10984338), DCX anti mouse 1:500 (Millipore Cat# AB2253, RRID:AB_1586992) VCP anti rabbit 1:2000 (Abcam Cat# ab11433, RRID:AB_298039), Iba1 anti rabbit 1:500 (Abcam Cat# ab178846, RRID:AB_2636859) TGF-Beta1 anti rabbit 1:1000 (Proteintech Cat# 21898-1-AP, RRID:AB_2811115).

## Results

### Eyeblink Conditioning

#### Microdrive

Microdrive implanted male, intact female, and ovx mice acquired tEBC over the course of 10 training sessions [F(1.819, 65.47) = 24.98, p= <0.0001] (Figure 2). Sex differences were observed in the rate of acquisition of tEBC in chronically implanted mice [F(2,36) = 6.941, p= 0.0028] (Table 1) (Figure 2). There was also a significant interaction between sex and session in chronically implanted mice [F(18, 324)=1.720, p=0.0348]. Male mice implanted with a chronic microdrive acquired tEBC significantly faster than intact female microdrive mice (Figure 2). However, ovariectomized females did not perform significantly differently from either intact female or male mice except for on T3 (Table 1) (Figure 2).

**Figure 2.**
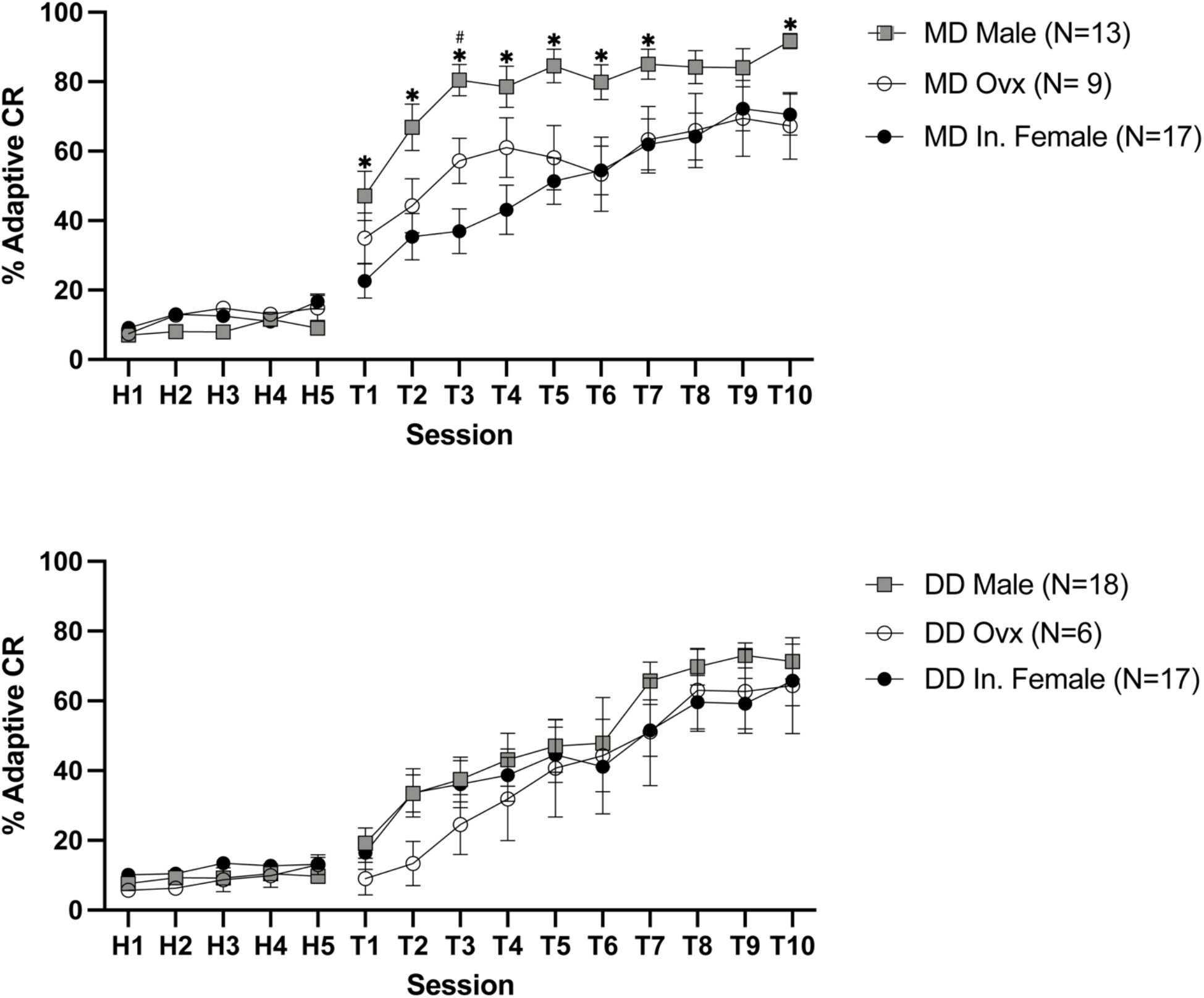
Learning Curve Comparison by Surgical Group. Data represents mean ± SEM **Upper**. Microdrive implant comparison by sex. * denotes p<0.05 MD Male to MD In. Female. # denotes p<0.05 MD Male to MD Ovx. **Lower.** Dummy Drive learning curve comparison by sex.

Number of trials to reach 8 consecutive CRs was also used as a measurement of learning rate. Animals that failed to reach 8 consecutive CRs by the end of 10 training sessions were scored as 500 trials, the total number of conditioning trials. Male microdrive mice reached 8 consecutive CRs significantly faster than intact female male [F(2, 36) = 7.985, p= 0.0013, Tukey’s Multiple Comparisons Test, Microdrive Male vs. Microdrive Female (p= 0.0012)] There were no significant differences between ovariectomized females and intact females or males (Table 1) (Figure 3). There was also a significant difference between the sexes in onset time of the CR of microdrive implanted animals [F(2, 36) = 4.480, p=0.0183] (Figure 4). Microdrive implanted males had an earlier onset compared to intact and ovariectomized female microdrive animals across multiple sessions (Table 1). However, onset time of the CR did not vary significantly across sessions (Table 1).

**Figure 3.**
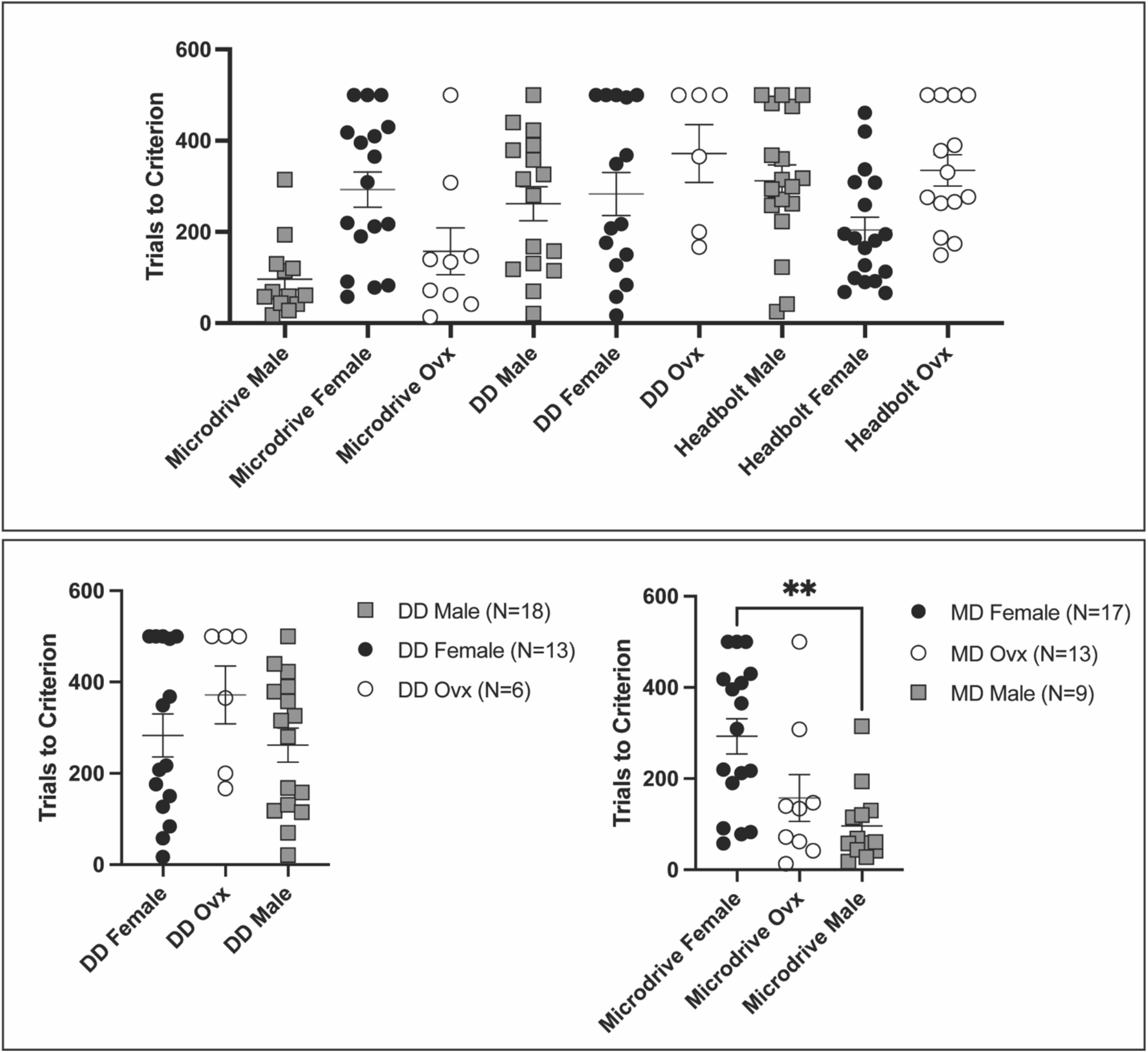
Trials to 8 Consecutive CRs Data represents mean ± SEM. **=p ≤ 0.01. **Upper**. Trials to 8 Consecutive CRs comparison by sex and surgical group. **Lower Left.** Dummy Drive comparison. No significant differences across sex. **Lower Right**. Microdrive comparison. Microdrive males take significantly fewer trials to reach 8 consecutive CRs, p=0.0275.

**Figure 4.**
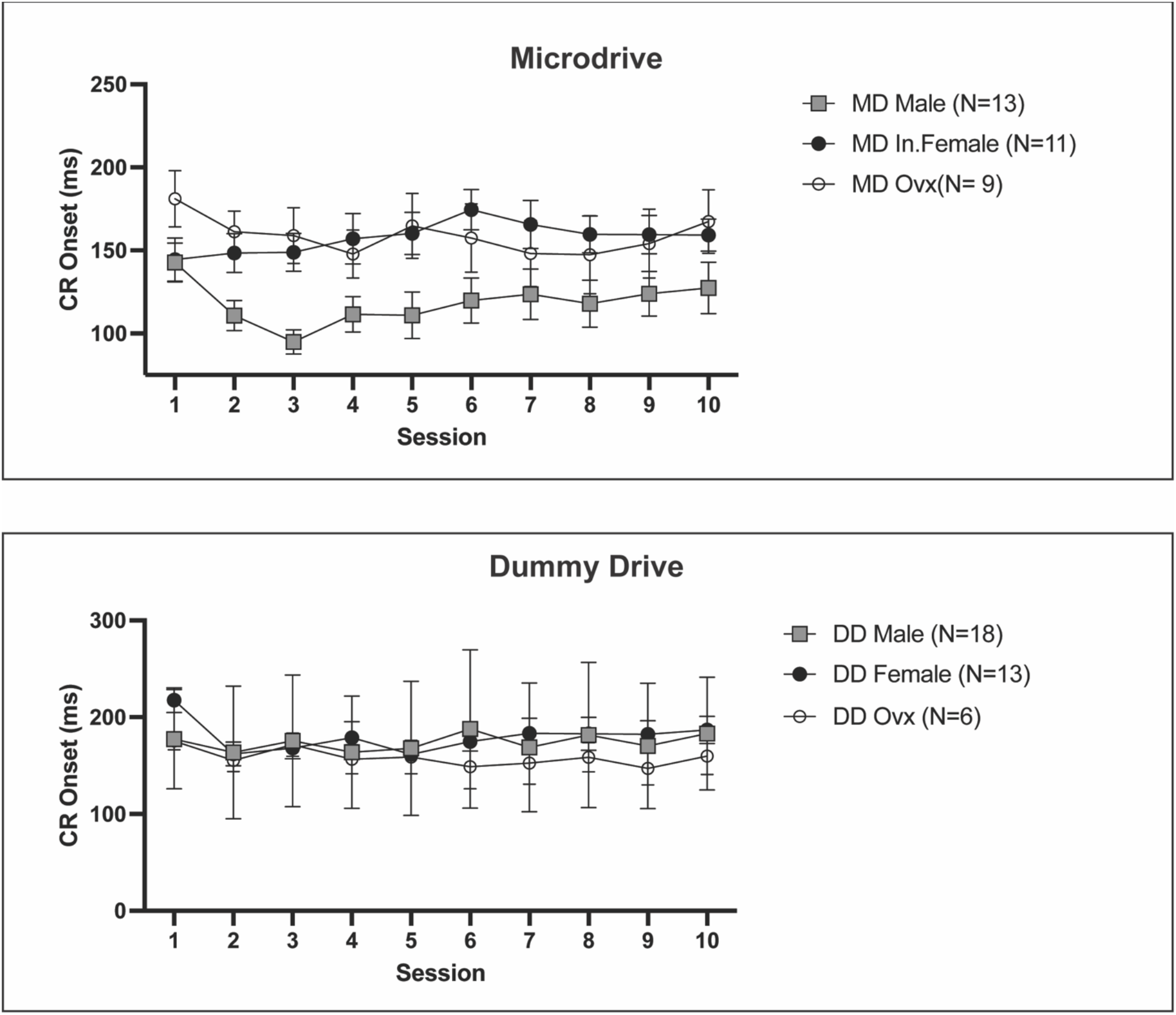
CR Onset Comparison of Dummy Drive and Microdrive. Data represents mean ± SEM **Upper.** Microdrive comparison of CR Onset. Microdrive males have a significantly lower CR onset, p= 0.0026. **Lower.** Dummy Drives show no significant differences in CR onset across sex.

#### Dummy Drive

The weight of the implanted microdrive alone did not lead to sex differences in acquisition. All animals with dummy drive implants acquired tEBC at similar rates (F(2, 38) = 0.5813, p=0.5640) (Figure 2). Additionally, across all groups, dummy drive animals were able to acquire tEBC over the ten sessions (F(1.554, 59.07) = 37.68, p=<0.0001). There were no significant differences between the sexes in trials to reach 8 consecutive CRs (Figure 3). Sex or session did not significantly affect the onset time of CRs (Table 1) (Figure 4).

#### Males

Male mice acquired tEBC at significantly different rates depending on the surgical procedure they received (F(2, 46) = 13.02, p=<0.0001) (Table 1) (Figure 5). Male microdrive mice learned tEBC significantly faster than both dummy drive and headbolt implanted males (Table 1). Across all surgical groups, male mice acquired tEBC (F (1.886, 86.77) = 57.84, p=<0.0001) (Figure 5). There was a significant interaction between surgical procedure group and session (F(18,414) = 3.280, p=<0.0001).

**Figure 5.**
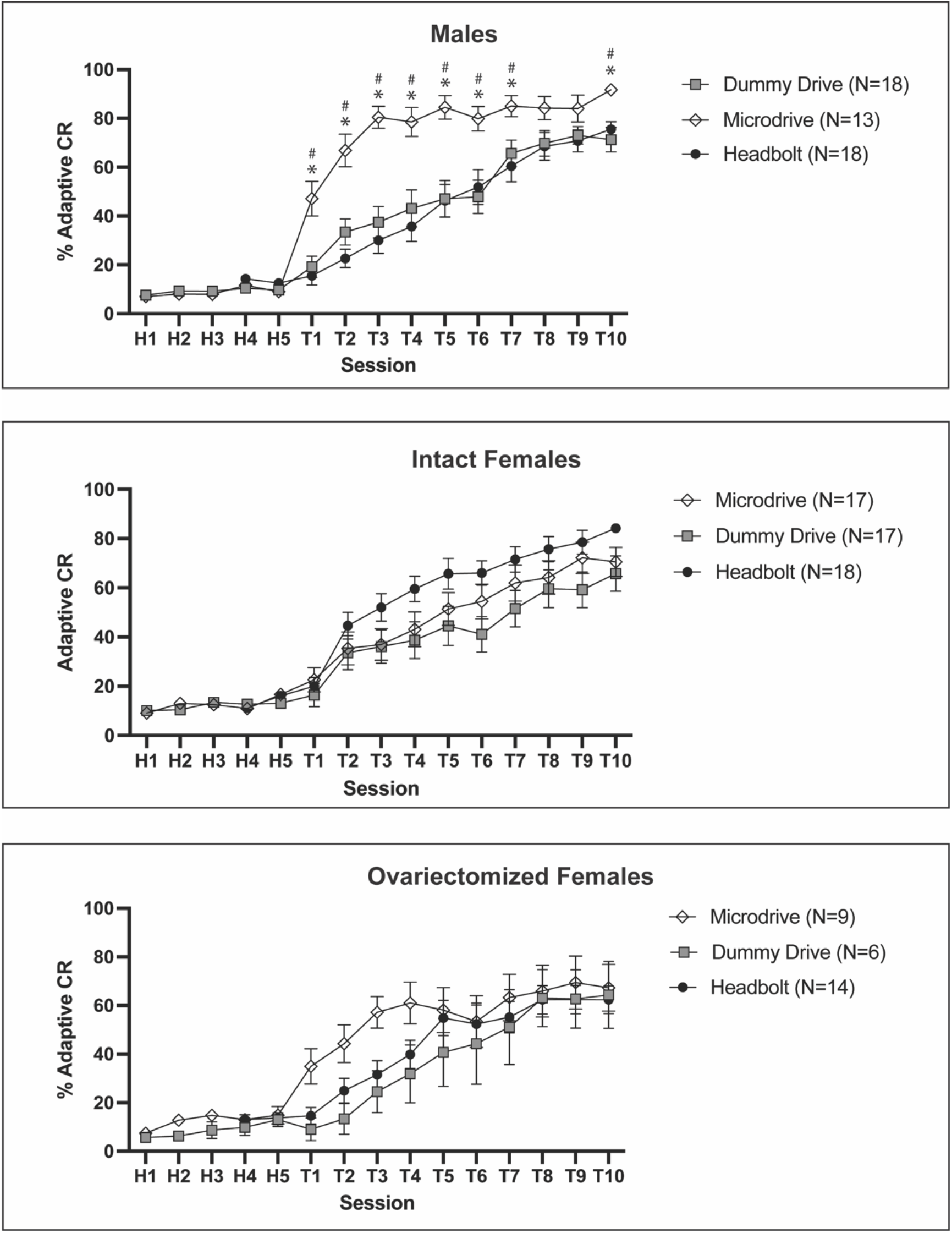
Adaptive CR Learning Curve Comparison. Data represents mean ± SEM **Top**. Male Comparison by Surgical Group. * denotes p<0.05 MD to HB, # denotes p<0.05 MD to DD. Microdrive males learned significantly faster than dummy drive and headbolt implanted males p<0.0001. **Center.** Intact Females CR Learning curves. **Bottom** Ovariectomized Female adaptive CR comparison by surgical group

Surgical procedure also had a significant effect on the number of trials male mice required to reach 8 consecutive CRs (F(2, 44) = 10.26, p=0.0002)(Figure 4). Male microdrive animals reached 8 consecutive CRs in significantly fewer trials compared to dummy drive males and headbolt males (Tukey’s Multiple Comparisons Test, Microdrive Male vs. DD Male p=0.0052, Microdrive Male vs. Headbolt Male, p=0.0002) (Figure 4). The effect of surgical procedure on tEBC was also observed through onset time of the conditioned response (F(2,44) = 6.842, p=0.0026) (Figure 7). Microdrive males have an earlier CR onset time compared to both dummy drive and headbolt implanted animals.

**Figure 6.**
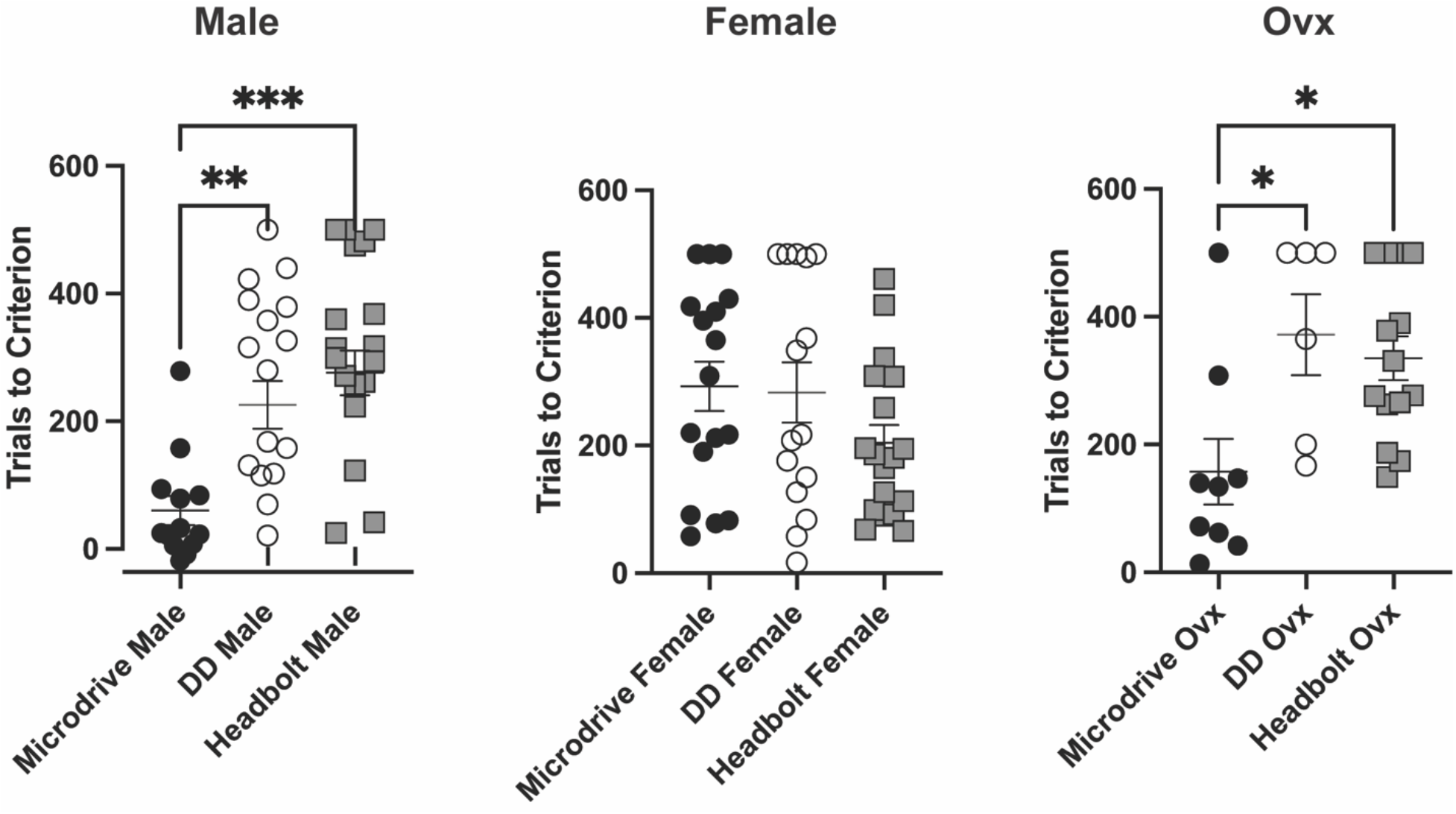
Trials to 8 Consecutive CRs Comparison by Sex. Error bars represent mean ± SEM *= p≤ 0.05, **=p ≤ 0.01, ***=p ≤ 0.001. **Left.** Male Trials to Criterion. Microdrive Males take significantly fewer trials to reach 8 consecutive CRs, compared to dummy drive and headbolt males, p=0.0448. **Center**. Intact Female Comparison of 8 consecutive CRs, no significant differences between groups. **Right.** Ovariectomized female comparison of 8 consecutive CRs. Microdrive ovx require significantly fewer trials to reach 8 consecutive CRs compared to dummy drive and headbolt ovx, p=0.0095.

**Figure 7.**
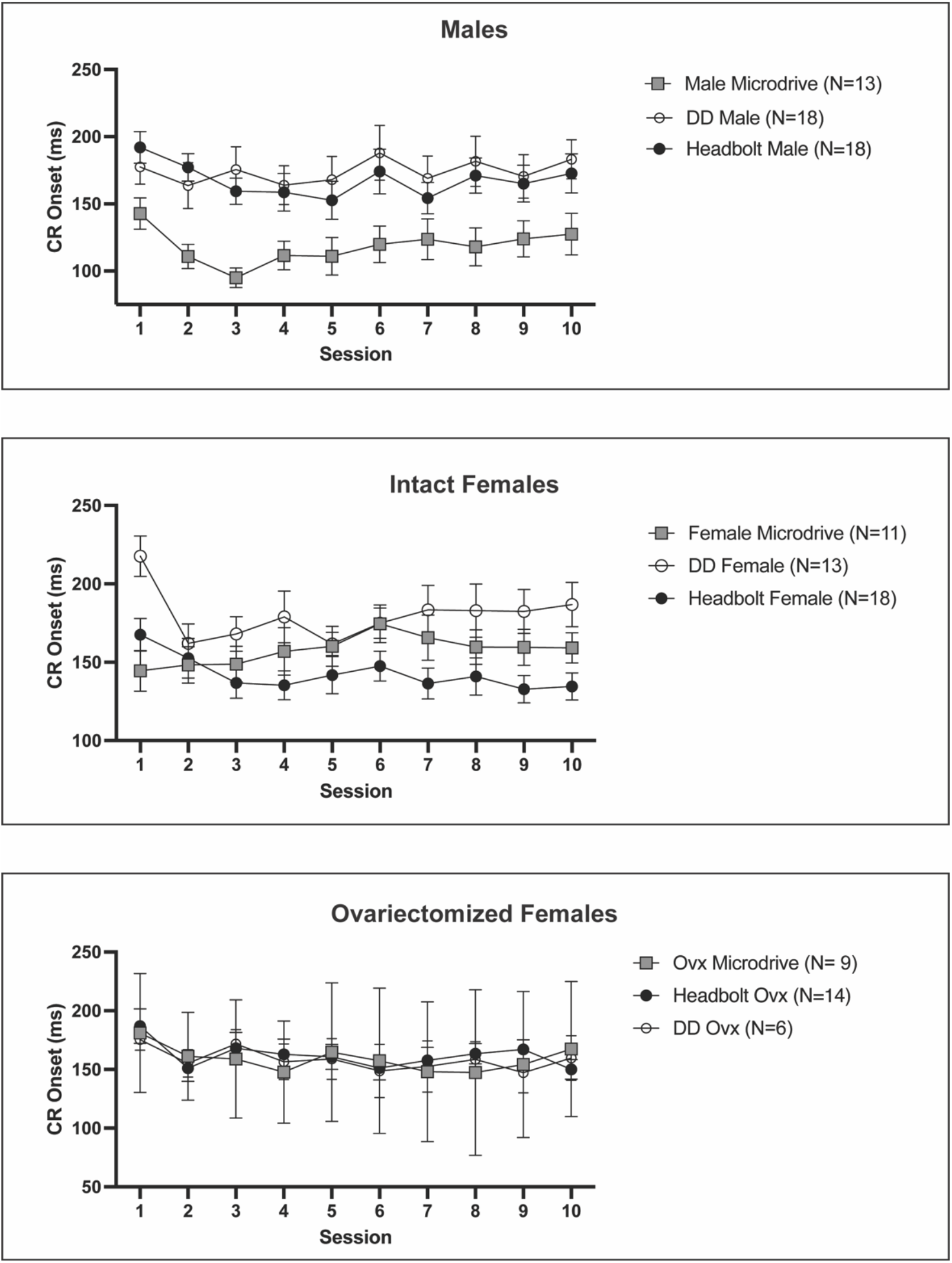
CR Onset Comparison by Sex. Data represents mean ± SEM **Top.** CR onset comparison in Males. Microdrive males have a significantly lower CR onset compared to male dummy drive and headbolt animals, p=0.0026. **Center.** CR onset comparison in Intact Females. Headbolt females had a significantly lower CR onset compared to microdrive and dummy drive females, p=0.0138 **Bottom.** CR onset in Ovx. No significant difference in CR onset in ovariectomized females.

#### Intact Females

There is a trend towards significance of surgical procedure on acquisition of tEBC in intact females (F(2, 49) = 2.877, p=0.0659) (Figure 5). Intact female microdrive and dummy drive animals tended to learn more slowly compared to headbolt implanted animals (Figure 5). Across all surgical groups, animals acquired tEBC (F(1.599, 78.35) = 58.61, p=<0.0001).

Surgical group did not significantly affect the number of trials required for intact females to reach 8 consecutive CRs (Table 1) (Figure 4). However, surgical procedure significantly impacted the timing of the CR of intact females (F(2, 47) = 4.696, p=0.0138) (Figure 7). There was also an interaction between surgical group and session on CR onset across all intact female groups (F(18, 422) = 1.688, p=0.0385).

#### Ovariectomized Females

Surgical group did not significantly affect the acquisition of adaptive CRs in tEBC in ovariectomized female mice (Table 1) (Figure 5). Across all groups, animals learned tEBC by the final session (F(1.711,44.49)=24.20, p=<0.001). However, there was a significant effect of the surgical group on the number of trials to reach 8 consecutive CRs (F(2, 26) = 5.604, p=0.0095) (Figure 6). Ovariectomized female microdrive mice required significantly fewer trials to reach 8 consecutive CRs (Table 1). There was no significant difference between surgical group or session on onset of the CR (Table 1) (Figure 7).

### Corticosterone Levels

#### All Subjects

To investigate if behavior differences seen among the sexes or surgery groups was due to corticosteroid level we performed a corticosterone ELISA assay. Across all animals, there was a significant effect of session on corticosterone levels (F(1.837,62.47)=3.64, p=0.0355). Significantly higher corticosterone levels were found in fecal matter collected during habituation week as compared to those from fecal matter collected from cages during the second week of training (Tukey’s Multiple Comparisons test, Hab vs T2, p=0.0385) (Figure 8).

**Figure 8.**
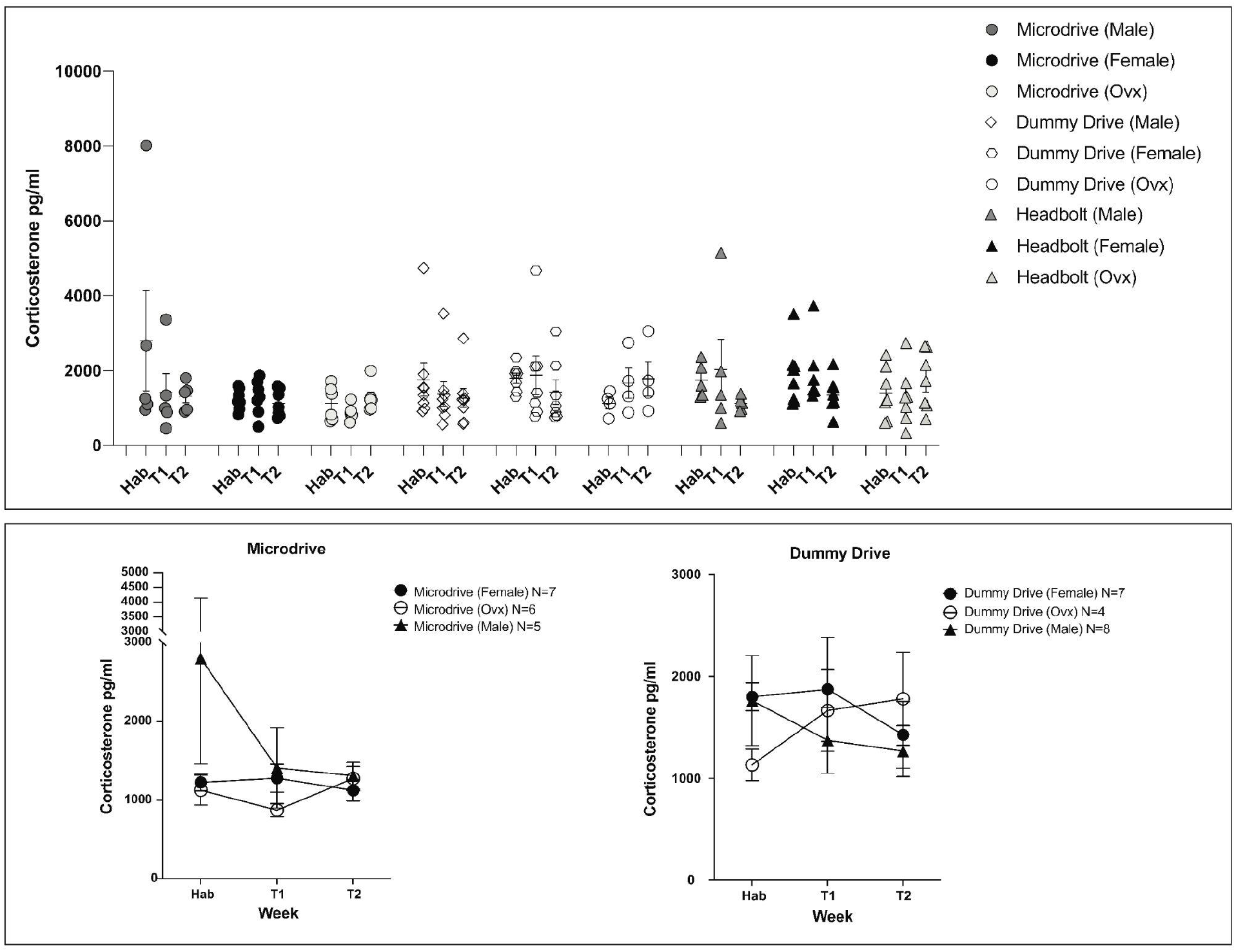
Corticosterone ELISA Analysis. Data represents Data represents mean ± SEM **Upper**. Corticosterone levels across all sex and surgical groups. **Lower Left**. Microdrive corticosterone levels across sex. **Lower Right.** Dummy Drive corticosterone levels across sex.

#### Microdrive/ Dummy Drive

Sex and session did not have a significant effect on corticosterone levels of microdrive or dummy drive animals (Table 1) (Figure 8).

#### Male/Intact Female/Ovariectomized Female

There was no significant effect of surgical group or session on corticosterone levels of male mice, intact female or ovariectomized female mice (Table 1) (Supplemental Figure 1).

### Tandem Mass Taq Spectrometry

To investigate potential mechanisms underlying sex differences in acquisition with chronic microdrive implantation, we performed a quantitative bottom-up proteomic screen using three 10-plex TMT-MS analyses (male, intact female, ovx). We set up the TMT analyses to compare protein abundance between surgical groups (headbolt, dummy drive, microdrive) for each sex (n=3 per group). We also utilized a float channel that was the same among all three TMT analyses, thus allowing us to compare across multiple TMT-MS screens and enabling comparison of sex within each surgery group (male, intact female, ovariectomized female). Overall TMT peak intensities and reporter ion intensities were similar across all TMT channels in each of the three TMT analysis groups (Supplemental Figure 2). The total number of quantified proteins were also similar across all three screens (Table 2).

We compared protein abundance in microdrive, dummy drive and headbolt mice per sex and identified significantly altered proteins (Table 2) (Supplemental Figure 3). To mine significantly altered proteins, we performed a GO: Biological Process Complete (GO:BPC) analysis for statistical overrepresentation with PANTHER.

Across the male dataset, significantly enriched GO:BPC terms included long-term synaptic potentiation, BDNF pathway, and positive regulation of MAPK pathway. To home in on the mechanisms underlying the sexually dimorphic enhancement of acquisition with chronic implantation, we further analyzed the significantly altered proteins between male microdrive and male dummy drive animals, as well as the comparison between microdrive and headbolt animals (Figure 9). Proteins significantly elevated in microdrive males compared to dummy drive males were enriched for several GO terms including: tumor necrosis factor-mediated signaling pathway, peroxisome organization and negative regulation of sodium ion transmembrane transporter activity (Figure 9). Meanwhile, proteins significantly decreased in microdrive compared to dummy drive mice were enriched for GO terms including: regulation of AMPA glutamate receptor clustering and NMDA selective glutamate receptor signaling pathway. Across the intact female dataset, significantly enriched GO:BPC terms included interleukin-1 signaling pathway, activation of MAPKK activity, and long-term synaptic potentiation. Further analysis showed that proteins significantly decreased in microdrive intact females compared to headbolt females were enriched in GO terms associated with the Protein Kinase A (PKA) pathway (Figure 10). Furthermore, proteins significantly decreased in microdrive intact females compared to headbolt intact females were enriched in GO terms associated with memory and inactivation of MAPK between microdrive and headbolt intact females (Figure 10). Finally, in the ovariectomized female group, significantly enriched GO:BPC term included: regulation of interleukin-6 production, dendritic spine development and regulation of MAP kinase activity. Compared to headbolt animals, ovariectomized microdrive females proteins with a significant decrease were enriched in GO terms associated with positive regulation of interleukin-6 mediated signaling pathway and negative regulation of corticotropin secretion (Figure 11). In addition, proteins significantly elevated in ovariectomized microdrive females compared to headbolt animals were significantly enriched in GO terms associated with dendritic spine development and regulation of AMPA receptor activity compared to headbolt animals. We also used the float channel in the TMT screen to compare normalized levels of proteins across sex/surgical group, in order to better understand what proteins and pathways may be involved in the observed sex differences in acquisition of tEBC. Among the most significantly altered proteins across sexes within each surgery group were: Mif, F3, Dusp7 and Nrtk2.

**Figure 9.**
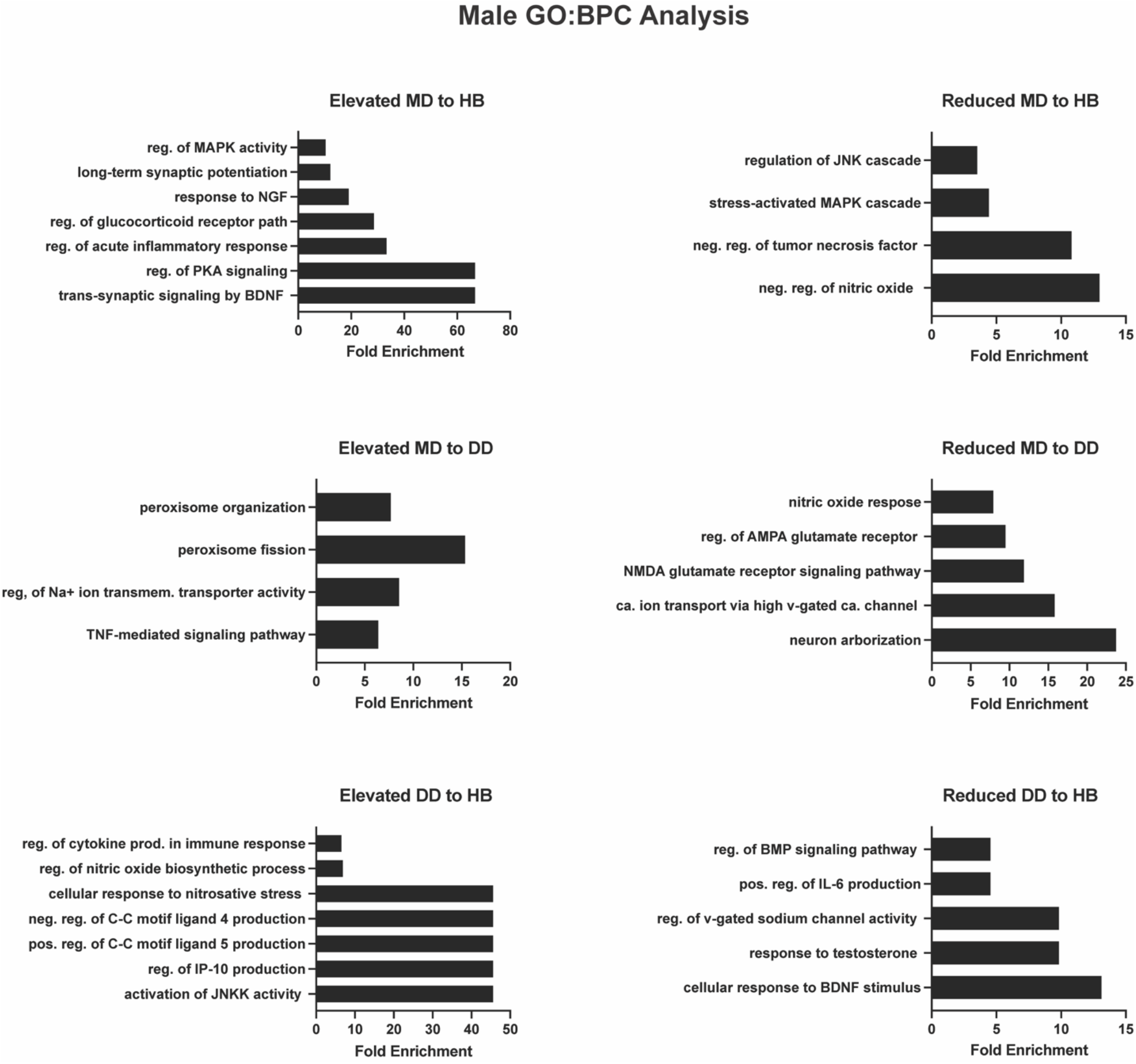
Male Panther GO:BPC Analysis. Gene ontology (GO) Biological Process Complete (GO:BPC) analysis of significantly altered proteins in male mice across microdrive, dummy drive and headbolt comparisons.

**Figure 10.**
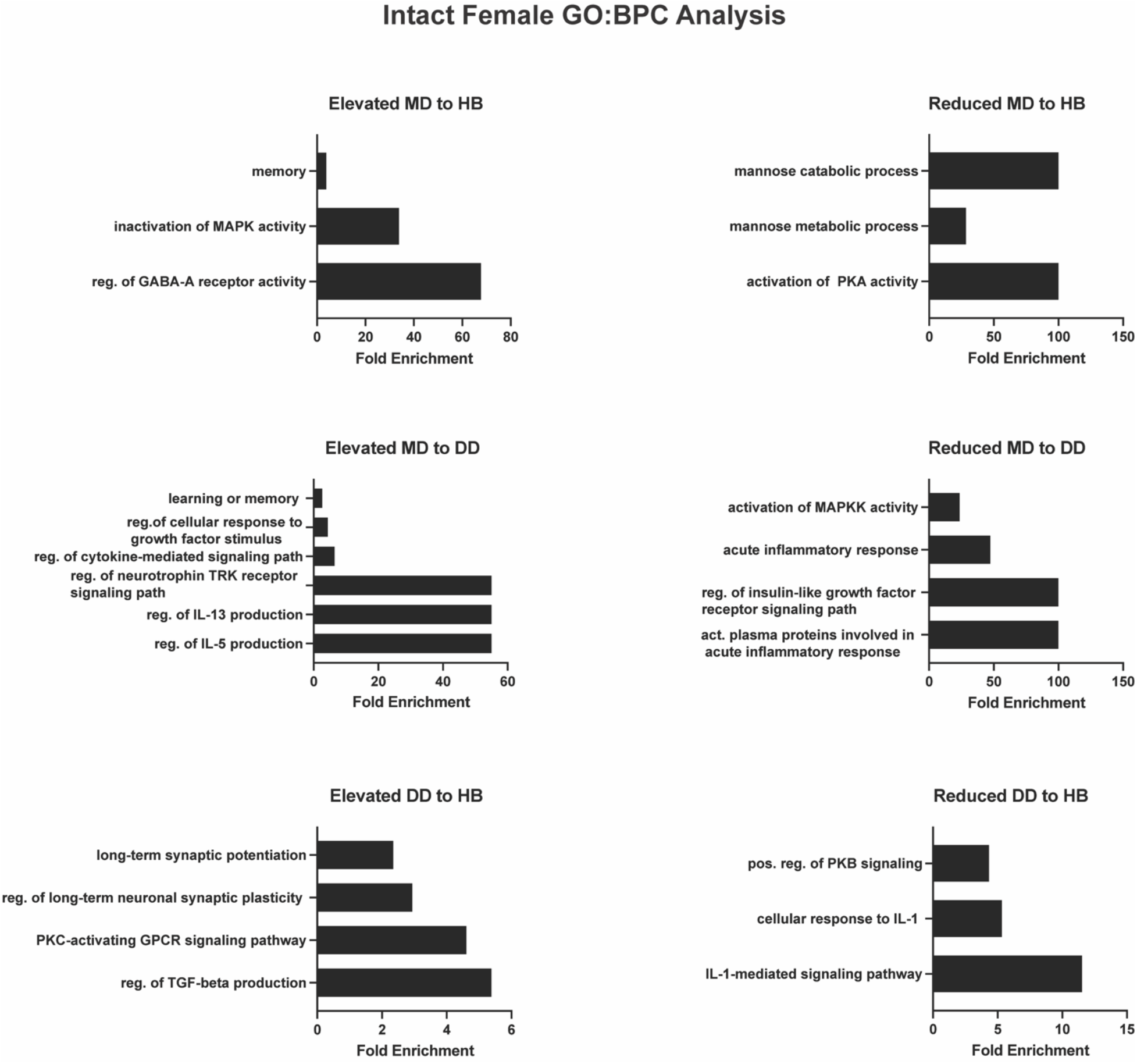
Intact Female GO:BPC Analysis. Gene ontology (GO) Biological Process Complete (GO:BPC) analysis of significantly altered proteins in intact female mice across microdrive, dummy drive and headbolt comparisons.

**Figure 11.**
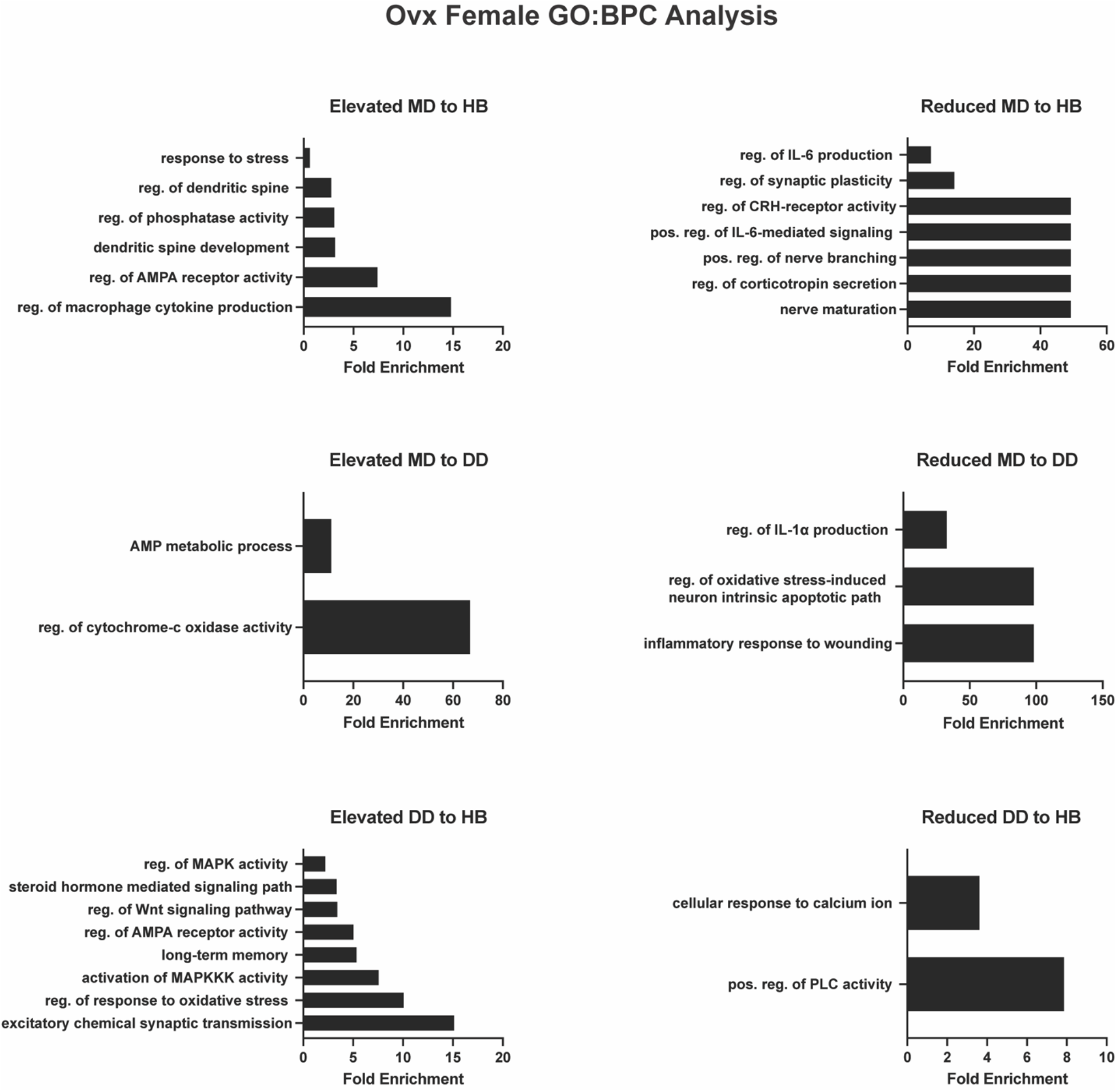
Ovariectomized Female GO:BPC Analysis. Gene ontology (GO) Biological Process Complete (GO:BPC) analysis of significantly altered proteins in ovariectomized female mice across microdrive, dummy drive and headbolt comparisons.

Macrophage migration inhibitory factor (Mif) is associated with the MAPK Pathway, acute inflammation, and Brain Derived Neurotrophic Factor (BDNF) and was significantly higher in intact female headbolt mice compared to both male and ovariectomized female headbolt mice (Table 3) (Figure 12).

**Figure 12.**
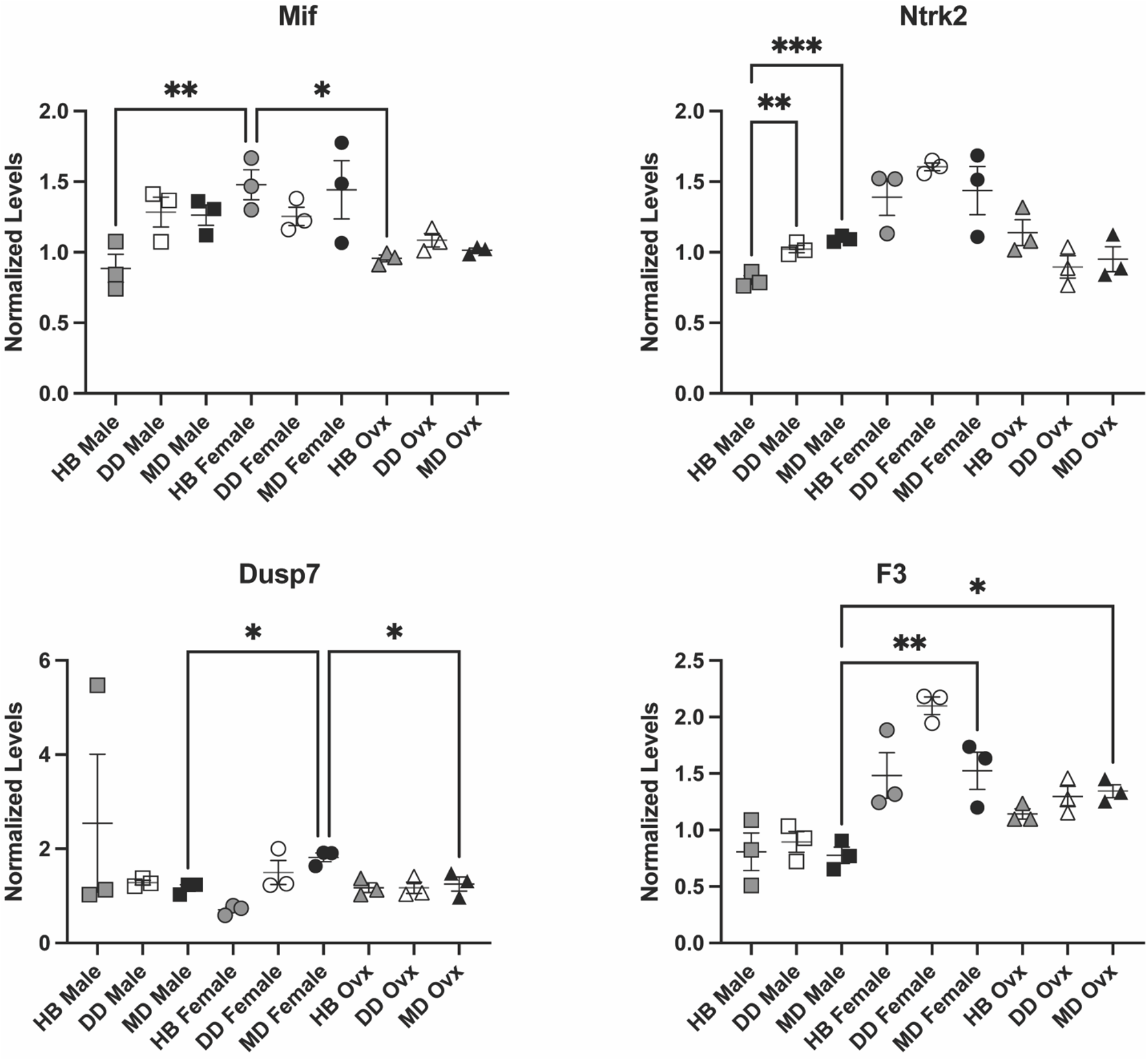
Normalized TMT-MS Protein Level Comparisons Across Groups. Data represents mean ± SEM. * = p≤ 0.05, **=p ≤ 0.01, ***=p ≤ 0.001. **Top Left.** Mif Normalized levels. Headbolt intact females had significantly higher levels of Mif compared to headbolt implanted males and ovx females, p=0.0411. **Bottom Left.** Dusp7 Normalized Levels across groups. Microdrive intact females had significantly higher levels compared to microdrive implanted males and ovx, p= 0.0109. **Top Right**. Ntrk2 normalized levels of protein across groups. Headbolt males had significantly lower levels compared to microdrive and dummy drive implanted males, p= 0.0003. **Bottom Right**. F3 Normalized levels of proteins across groups. Microdrive implanted males had significantly lower levels of F3 compared to intact and ovx females, p=0.0068.

Coagulation factor III (F3) is associated with activating plasma proteins involved in the acute inflammatory response. F3 levels were significantly lower in male microdrive animals compared to both intact and ovariectomized microdrive females (Table 3) (Figure 12). Male microdrive animals acquired tEBC faster than both intact and ovariectomized females.

Dual specificity protein phosphatase 7 (Dusp7) is associated with the MAPK pathway and was significantly higher in microdrive intact females compared to males and ovariectomized females with microdrive implants (Table 3) (Figure 12). Microdrive intact females learned significantly slower than microdrive males (Figure 2). Dusp7 levels were also significantly lower in intact female headbolt animals compared to both microdrive and dummy drive implanted female mice (Table 3) (Figure 12).

BDNF/NT-3 growth factors receptor (Ntrk2) is associated with a response to nerve growth factor and BDNF and was significantly lower in headbolt males compared to microdrive and dummy drive males (Table 3) (Figure 12). Proteins associated with the MAPK pathway, BDNF and acute inflammation were found to be significantly altered in our TMT-MS screen.

### Western Blot

Western Blot analysis did not show significant differences in neuroinflammatory or neurogenesis markers across sexes in the Dummy Drive and Microdrive groups (Table 4) (Supplemental Figure 4). Headbolt animals showed significant differences in levels of TGF-β (F(2,15)= 3.965, p=0.0415). Ovx females had significantly higher levels of TGF-β compared to intact females, (Tukey’s Multiple Comparisons Test, p= 0.0398). No other markers showed significant differences across sexes (Table 4).

When comparing across surgical groups, males, intact females and ovariectomized females showed no significant differences in levels of GFAP (Table 4) (Figure 13). Males also did not show significant differences across surgical groups in DCX or TGF-β (Table 4).

**Figure 13.**
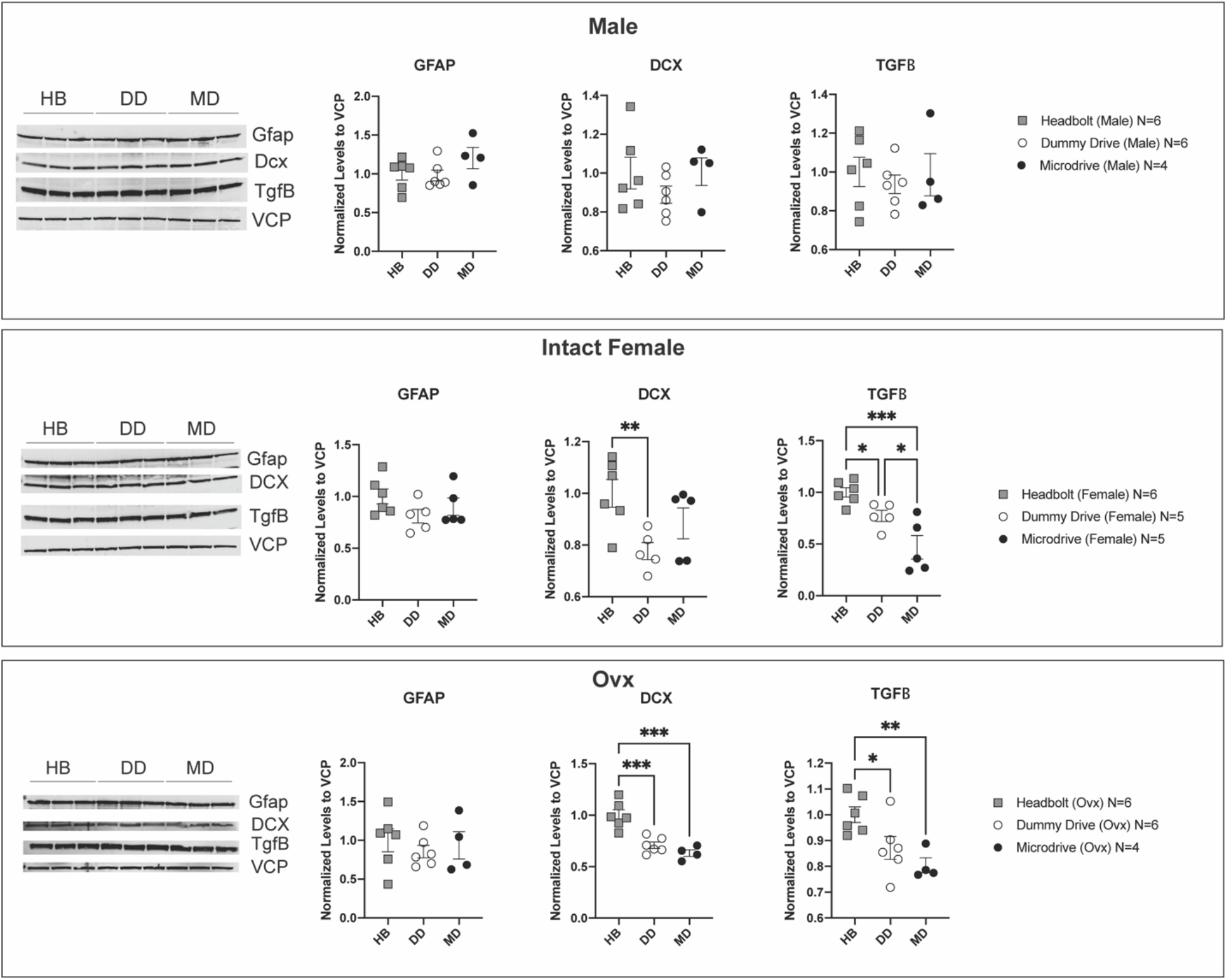
Western Blot Analysis by Sex. Data represents mean ± SEM. * = p≤ 0.05, **=p ≤ 0.01, ***=p ≤ 0.001. Top. Male Western Blot Analysis. Center. Intact Female Western Blot Analysis. Levels of DCX were significantly higher in headbolt intact females compared to dummy drive implanted females, p= 0.025. Levels of TGFβ were significantly lower in microdrive females compared to headbolt and dummy drive implanted females, p=0.0007. Bottom Ovx Western Blot Analysis. Levels of DCX were significantly higher in headbolt ovx compared to dummy drive and microdrive implanted ovx, p=<0.0001. Levels of TGFβ were significantly lower in microdrive ovx compared to headbolt and dummy drive implanted ovx, p=0.0094.

However, both intact females and ovariectomized females showed significant differences in TGF-β levels across surgical groups [Intact Female: F(2,13)= 13.31,p=0.0007, Ovx: F(2,13)= 6.837,p=0.0094](Figure 13). Levels of TGF-β in intact females decreased between headbolt and microdrive mice as well as dummy drive and microdrive mice (Table 4). TGF-β levels in ovariectomized females were significantly lower in microdrive compared to headbolt animals. (Table 4).

Intact and ovariectomized females also showed significant differences in DCX between surgical groups (Table 4). Whereas intact female DCX levels decreased with chronic dummy drive implant, ovariectomized female DCX levels decreased with both microdrive and dummy drive implantation (Table 4).

## Discussion

Sexually dimorphic behaviors were observed in tEBC acquisition with surgical implants. Female mice with headbolts alone learned faster than male and ovariectomized females. However, implantation of a dummy drive or microdrive on female mice resulted in a delay in learning compared to headbolt females (Figure 2). In contrast, microdrive implanted males acquired tEBC at a significantly faster rate compared to dummy drive and headbolt implanted males (Figure 3). Additionally, microdrive implanted males learned faster than microdrive implanted females (Figure 3).

Sexually dimorphic behaviors have previously been observed in stressed animals, therefore we investigated whether the surgeries and implantations led to an increased stress response and increased concentration of corticosterone (Wentworth-Eidsaune, Hennessy, and Claflin 2016; Wood and Shors 1998). Corticosterone analysis from fecal matter collected from all animals showed a significant decrease in corticosterone from habituation week to training week two, suggesting that animals have the highest stress levels following surgery and then acclimate to the head-fixed apparatus during the conditioning period. However, there were no significant differences observed between surgical groups or sexes, suggesting different surgeries, of particular note, the microdrive surgery where tetrodes are implanted into the cortex, do not affect corticosteroid levels. As our analysis grouped fecal matter over the course of a week, it is plausible that sex and/or group differences had inadequate signal-to-noise. In addition, it is possible that surgery and tEBC alone leads to increased corticosterone that is not detectably increased by additional stress of the microdrive implant. Notably, Shors et al. similarly found no significant differences in corticosterone levels between acutely stressed and conditioned rats when trunk blood was collected for corticosterone serum analysis (Shors, Weiss, and Thompson 1992).

Among the learning differences seen between groups was the result that microdrive males learned significantly faster than males in the other surgery groups. Since dummy drive males also received chronic implants, stress due to the weight of the implant likely does not account for facilitation. Rather, these differences in acquisition suggest an additional mechanism of facilitation due to additional surgical procedures including craniotomy and implanted tetrodes. These aspects of microdrive implantation may lead to a neuroinflammatory response (Patrick et al. 2011; Polikov, Tresco, and Reichert 2005; Prasad et al. 2012) that supports learning in males but not intact or ovariectomized females. Therefore, we performed Western Blot analysis to investigate possible neuroinflammatory mechanisms underlying the differences observed between groups. We also performed TMT-MS to investigate what proteins or protein pathways were significantly altered among the sexes and/or the surgery groups.

Surgical implantation of the microdrive may serve as a neuroinflammatory priming event for male mice, leading to facilitation during subsequent stressful events, including acquisition of eyeblink conditioning (Frank et al. 2007). When male rats received chronic restraint stress and subsequent lipopolysaccharide (LPS) endotoxin injection, they showed exaggerated levels of proinflammatory cytokines IL-1β and TNF-α compared to primed females (Bekhbat and Neigh 2018; Munhoz et al. 2006). In addition, proinflammatory cytokines have been shown to facilitate acquisition of water maze (Gibertini 1998), delay eyeblink conditioning in Sprague-Dawley rats(Servatius and Beck 2003), and recall of fear conditioning (Goshen et al. 2007). Indeed, our TMT-MS GO:BPC analysis showed that proteins associated with the TNF-α pathway were significantly altered in male animals (Figure 9). Therefore, neuroinflammatory priming due to the implantation of tetrodes may underlie observed sexually divergent behavior in acquisition.

Activation of the “alternative” neuroinflammation pathway could also underlie facilitation in acquisition in male microdrive mice. Alternatively activated microglia express anti-inflammatory cytokines interleukin-10 (IL-10), transforming growth factor-β (TGF-β), as well as growth factors including NGF and brain derived neural growth factor (BDNF) (Kohman and Rhodes, 2013). Activation of the “alternative” pathway promotes neuroprotective and regenerative processes (Colton 2009; Kohman and Rhodes 2013). TMT-MS analysis showed significant changes in both Mif and Nrtk2 levels in male mice. Mif and Nrtk2 are associated with BDNF, indicating that the alternative pathway may also play a role in tEBC acquisition. Together, activation of enhanced neurogenesis through activation of the alternative pathway and increase of proinflammatory cytokines like TNF-α may lead to the facilitation of tEBC in microdrive males. Future investigation into proinflammatory cytokines such as IL-1β and TNF-α, may further elucidate the underlying mechanisms underlying sexually dimorphic facilitation.

Microdrive intact females are impaired in acquisition of tEBC compared to males, due to changes in the MAPK pathway and neurogenesis, as suggested by TMT-MS and Western Blot analysis (Figure 2). In particular, Dusp7 and Mif, which are associated with the MAPK pathway, were among the significantly altered proteins in intact female microdrive mice (Figure 12). Inactivation of the MAPK pathway has been shown to impair learning and LTP(Philips et al. 2013; Schafe et al. 2000; Atkins et al. 1998). In addition, the impairment of learning fear conditioning was dose dependent, suggesting that even small amounts of MAPK inactivation may affect learning (Atkins et al. 1998).

Doublecortin is associated with levels of neurogenesis, which has a well-documented role in learning (Miller, Weiss, and Disterhoft 2019; Dalla et al. 2009; Deng, Aimone, and Gage 2010). Levels of doublecortin were significantly lower in intact microdrive females and dummy drive females compared to headbolt females. This decrease in neurogenesis may underlie the observed behavioral differences and may be further connected to the altered levels of MAPK (Waltereit and Weller 2003; Jiang et al. 2015). The decrease in neurogenesis was not dependent on sex hormones, as ovariectomized headbolt females also had higher levels of doublecortin compared to microdrive and dummy drive ovariectomized females (Figure 13). However, sex steroids including estrogen, progesterone and testosterone have known roles in the regulation of the immune system and provide further explanation for observed sex differences in acquisition (Berkiks et al. 2019). The influence of estradiol on the HPA axis may in part explain trends in differences between intact and ovariectomized microdrive females (Figure 2).

Neuroinflammatory and neurogenesis processes underlie the sexually dimorphic behavior observed with surgical implantation of the microdrive, where male microdrive animals learned more quickly than intact and ovariectomized females. Our TMT-MS analysis suggest the MAPK pathway, BDNF and acute inflammation may play a critical role in sex and surgical differences observed in acquisition. Observed sex differences in doublecortin and TNF-α also support the notion that neuroinflammatory and neurogenesis systems shape the differences in acquisition of eyeblink conditioning. Together, these conclusions emphasize the need to consider the behavioral effects of sex dependent changes in neuroinflammation and neurogenesis due to experimental manipulation.

## All Tables are Includes as Separate Tabs in the Attached Table Spreadsheet

**Table 1:** Statistical table for behavioral and corticosterone analysis. Includes all analysis for Adaptive CR, CR Onset, Consecutive 8 CRs.

**Table 2:** Number of significantly enhanced and reduced proteins per sex and surgical group.

**Table 3:** Statistical analysis for significantly altered proteins across sex and surgical group.

**Table 4:** Statistical data for Western Blot analysis.

## Declarations

### Funding

37 AG008796 (Disterhoft)

R01 NS113804 (Voss, Disterhoft)

