## Supplemental Figures 1-4 for "Sex-Dependent Effects of Chronic Microdrive Implantation on Acquisition of Trace Eyeblink Conditioning"

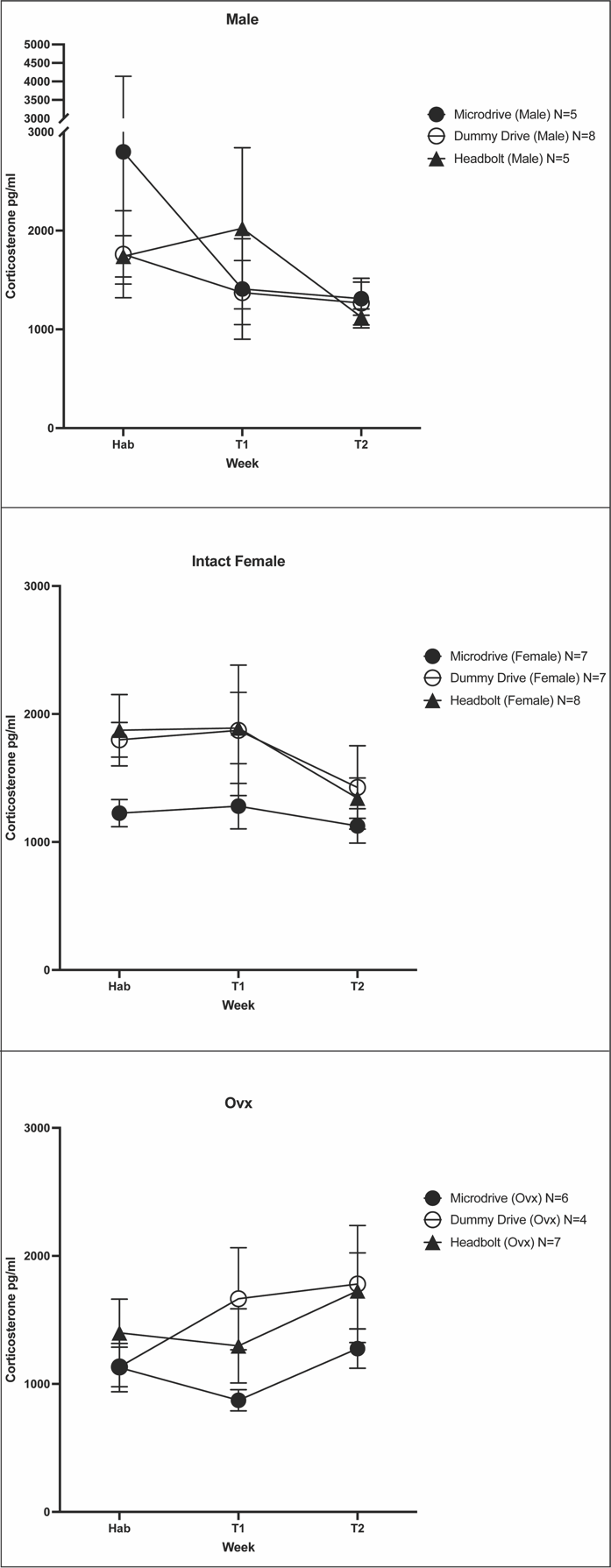

**Supplemental 1**. Corticosterone Analysis by Sex (pg/ml). **Top**. Male Corticosterone Analysis **Center**. Intact Female Corticosterone Analysis. **Bottom.** Ovx Corticosterone Analysis.

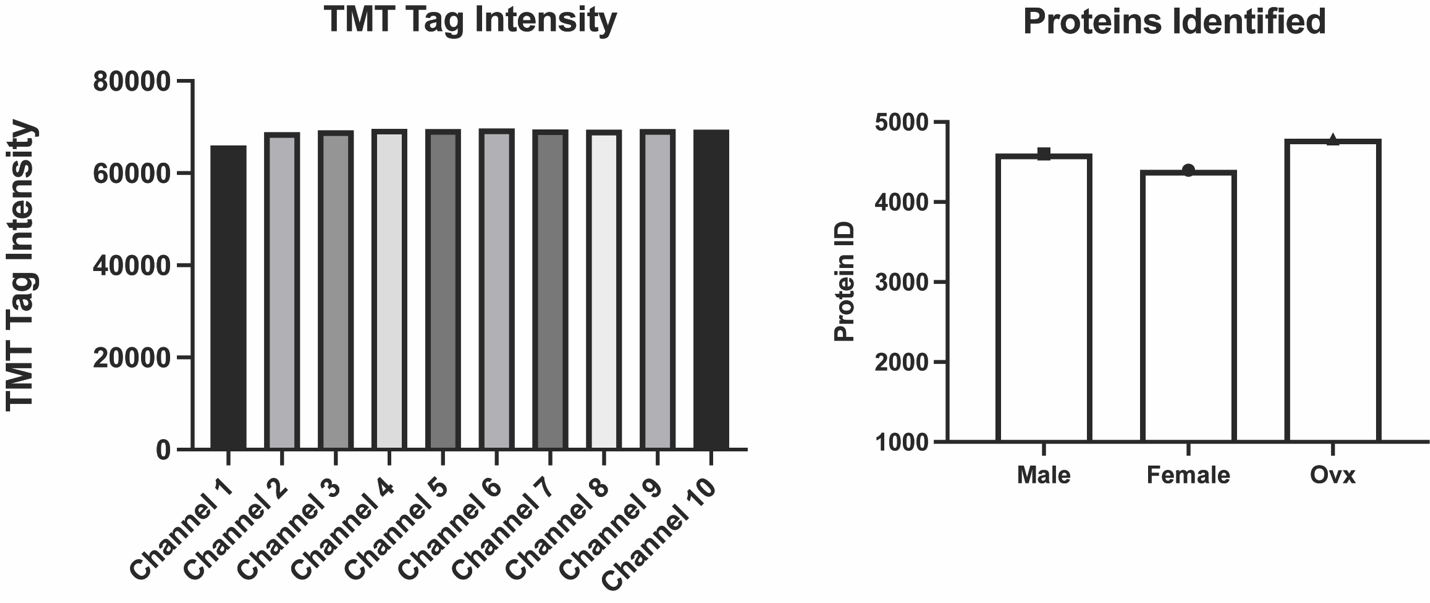

**Supplemental 2.** TMT-MS Tag Intensity and Number of Proteins Identified.

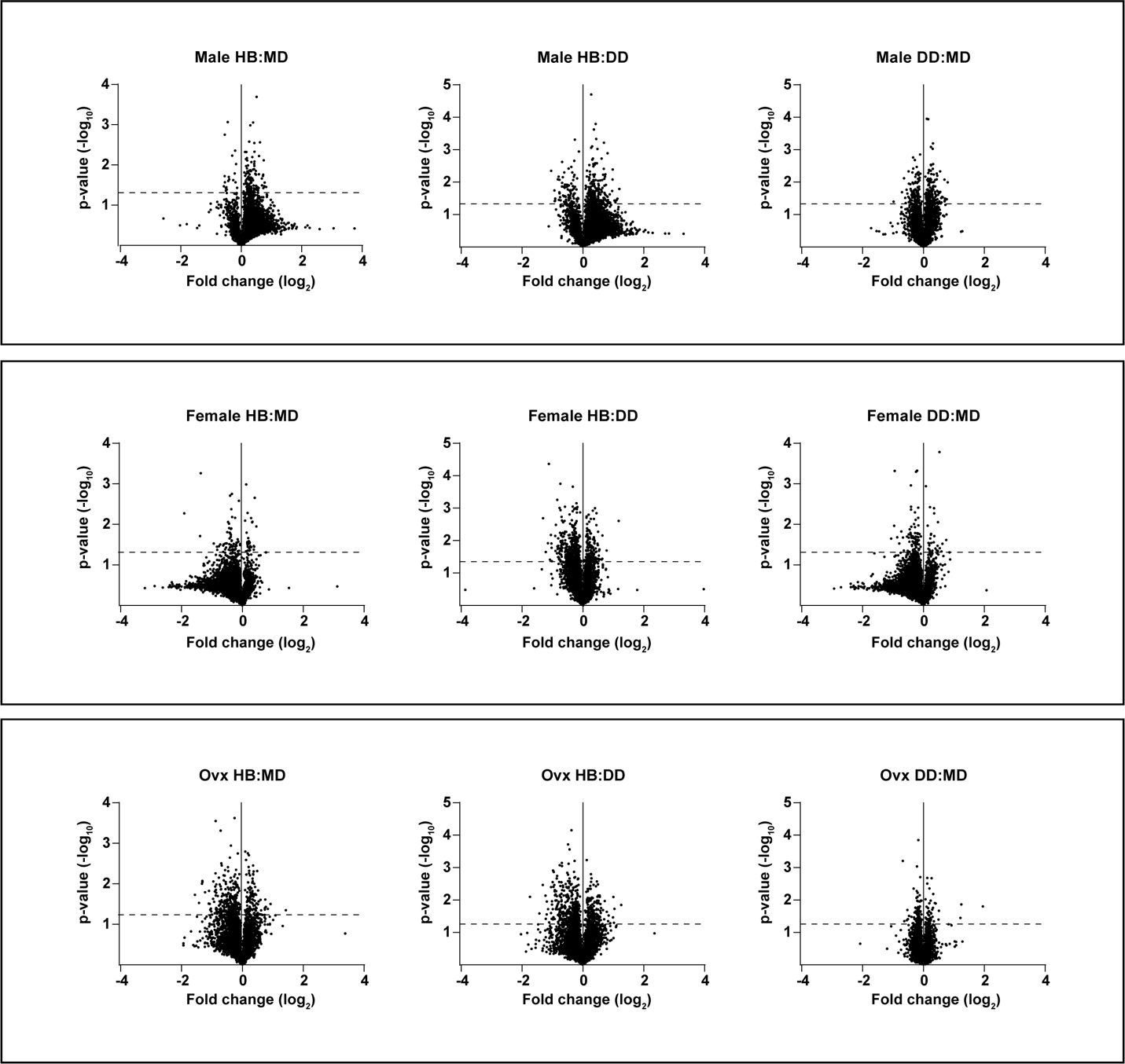

**Supplemental 3.** Volcano plots depicting protein fold change for paring listed in the title of the plot. Significant proteins (p-value < 0.05) are above the dotted line (Table 2).

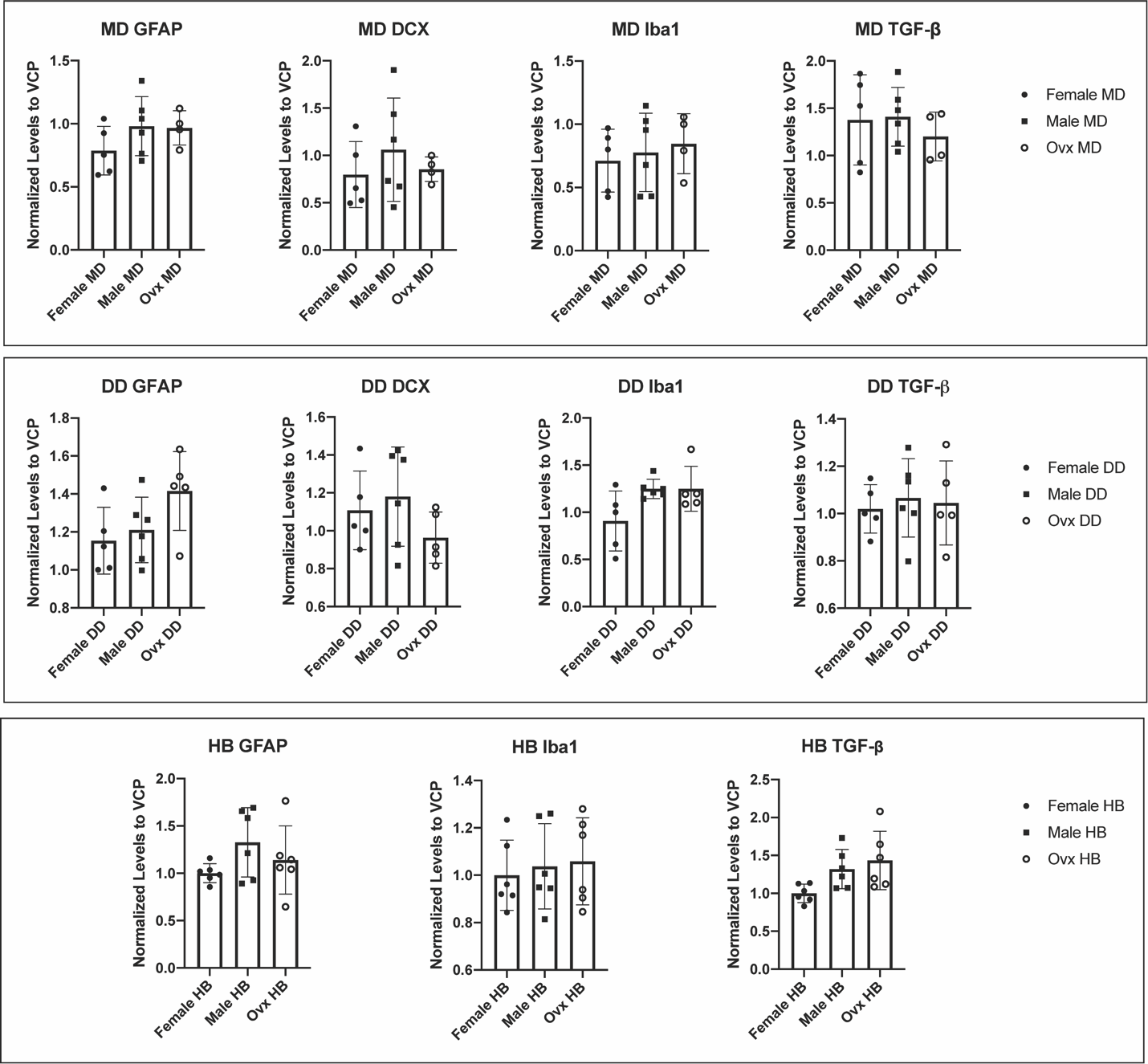

**Supplemental 4**. Western Blot Analysis by Surgical Group. Top. Microdrive Western Blot (Male MD=6, In. Female MD= 5,Ovx MD= 4) Center. Dummy Drive Western Blot Analysis (Male DD= 6, In. Female DD=5 Ovx DD= 5 ) Bottom. Headbolt Western Blot Analysis(Male HB= 6, In. Female HB=6 , Ovx HB=6)
